# Npl4 decodes polyubiquitin length and gates D1-D2 coupling in human VCP/p97

**DOI:** 10.64898/2026.05.01.722286

**Authors:** Laxmikanta Khamari, Jingxuan Tang, Stephanie L. Moon, Nils G. Walter

## Abstract

VCP/p97 binds the Npl4**–**Ufd1 heterodimer adaptor to extract polyubiquitinated substrates for proteasomal degradation, but how it decodes K48-linked chain length and how D1-coupled events license downstream D2 power strokes remain unclear. Here we introduce smUbiRAD, or single-molecule ubiquitin recognition and dynamics, and identify a sharp chain-length threshold: Npl4 binds transiently to short chains but switches to long-lived, multivalent engagement on tetra- and penta-ubiquitin. Ufd1 and p97 further stabilize these complexes mainly by suppressing Npl4 dissociation without affecting initial encounter. In fully assembled p97-Ufd1-Npl4-substrate complexes, D1 ATP hydrolysis—rather than D2—drives rapid Npl4 exchange. These results support a model in which D1-powered conformational changes promote cofactor Npl4, but not Ufd1, turnover and gate iterative coupling to downstream D2-driven substrate processing. Finally, we show that multisystem proteinopathy variants R155H and A232E bias p97 toward a high-affinity resting state and accelerate Npl4 exchange, implicating hyperactive cofactor cycling as a disease-linked dysregulation.

## Introduction

Cells rely on ATP-driven protein-remodeling machines to maintain protein homeostasis. The hexameric AAA+ ATPase VCP/p97 extracts ubiquitylated protein clients from membranes, ribonucleoprotein assemblies and macromolecular complexes and directs them toward proteasomal and autophagic clearance pathways^1–7^. Through this broad client spectrum, p97 safeguards endoplasmic-reticulum protein quality control, ribosome-associated quality control, chromatin-associated repair, and stress granule dynamics, placing it at the core of cellular proteostasis^1, 3, 8–11^. Missense mutations in p97 are associated with multisystem proteinopathy (MSP)—a dominantly inherited disorder encompassing inclusion-body myopathy, Paget disease of bone, frontotemporal dementia (FTD) and amyotrophic lateral sclerosis (ALS)—and are linked to cancer and cardiomyopathy^7, 12–15^. The R155H allele is the most common MSP-associated variant, and the A232E mutation is associated with severe disease^15–18^. Although such variants counterintuitively elevate ATPase and unwinding activity in bulk assays^19–21^, their single-turnover effects on ubiquitin decoding and cofactor exchange remain largely unresolved, limiting precision therapeutic strategies.

p97 assembles as a homohexamer with stacked D1 and D2 ATPase rings and flexible N-terminal domains (NTDs) that couple nucleotide state to cofactor binding and substrate processing^22, 23^. There are over 40 cofactors that have been identified to interact with p97, which aid VCP substrate identification and recruitment. The Npl4**–**Ufd1 complex is a highly characterized p97 cofactor that recognizes K48-linked ubiquitin chains and is implicated in diverse processes including ribosome-associated quality control, ER-associated degradation, chromatin-associated protein extraction, and stress granule disassembly^1, 2, 11, 14, 24–27^. Studies of Cdc48, the yeast homologue of p97, indicate that the C-terminal region of Npl4 binds polyubiquitin multivalently and stimulates ATPase activity in a linkage- and chain-length-dependent manner. Ufd1 engages a groove on Npl4, anchoring the heterodimer to the D1 ring and positioning the polyubiquitin chain above the central pore^21, 28–30^. Recent cryo-EM analyses of human p97-Npl4**–**Ufd1 complexes identified multiple “seesaw” conformations of Npl4 on D1, in which zinc-finger motifs alternately contact the NTDs and p97 pore, suggesting a dynamic mechanism for regulating substrate access and unfolding^28^.

Despite these structural insights, key questions remain. First, it is unclear whether polyubiquitin chain-length dependent decoding by human Npl4 is a static or dynamic process. The minimum ubiquitin chain length has not been quantitatively established, in part because prior studies used long chains or truncated Npl4. Second, while structural snapshots capture Npl4–Ufd1–p97 complexes in defined nucleotide states, how Npl4 senses chain length and dynamically couples it to ATP hydrolysis in the D1 and D2 rings to control cofactor turnover remains unresolved. Third, the respective roles of Npl4 and Ufd1—whether both exchange on substrate or whether one serves as a scaffold—remain poorly defined, particularly in relation to nucleotide-driven NTD “UP/DOWN” conformational switching^31^. NTD switching is disease relevant: cryo-EM and biophysical studies show that MSP-associated mutations such as R155H and A232E at the NTD–D1 interface destabilize the “DOWN” state and favor an ATP-like “UP” conformation, increasing ATPase activity^32–36^, yet how D1 timing relates to Npl4/Ufd1 residence is unknown.

Here, we developed a single-molecule ubiquitin recognition and dynamics (smUbiRAD) framework that quantifies how human Npl4, alone and in complex with Ufd1**–**p97, decodes K48-linked polyubiquitin chain length and dynamically connects this readout to ring-specific ATPase activity. Using SNAP-tagged model substrates bearing one to five ubiquitins, we combined bulk fluorescence polarization, multicolor total internal reflection fluorescence (TIRF) microscopy, ATPase assays, Walker A mutants, and MSP-linked VCP variants to measure affinities, kinetics, and nucleotide-dependent cofactor exchange. We identify a sharp Npl4 binding threshold at tetra- and penta-ubiquitin, show that Ufd1 and VCP primarily stabilize bound states by suppressing dissociation, and demonstrate that D1 ATP hydrolysis drives Npl4 exchange while Ufd1 remains p97 bound. Disease-associated mutants R155H and A232E tighten binding in the resting state, yet accelerate ATP-driven Npl4 turnover, consistent with detrimental hyperactive cofactor cycling. Together, these findings define a dynamic chain-length decoding mechanism and a D1-facilitated cycle of cofactor exchange that gates iterative D2-driven substrate processing. More broadly, this work shows how Npl4 integrates ubiquitin-chain architecture and ATPase-state control to govern human p97 function, sharpening mechanistic understanding of substrate selection and proteostasis.

## Results

### Nonlinear effects of the substrate’s polyubiquitin chain length on Npl4 selection

To systematically interrogate the impact of polyubiquitin chain length on cofactor engagement and p97 activation, we first established a robust workflow for the recombinant synthesis and purification of SNAP-tagged model substrates, each bearing precisely one to five K48-linked ubiquitin moieties (Ub^1^–S to Ub^5^–S; Fig. 1a, Supplementary Tables 1 and 2, and Supplementary Fig. 1). A SNAP-tag was fused to the C-terminus of ubiquitin and recombinantly expressed in *E. coli*, followed by purification (Methods Sections). Subsequent chain elongation was performed *in vitro* using enzymatic polyubiquitination (Fig. 1a and Supplementary Fig. 1) with ligase specific E1 and E2-3 (gp78RING-Ube2g2 chimera, Supplementary Tables 1 and 2, Supplementary Fig. 2)^14, 21^. SDS-PAGE analysis of both biotin- and Cy3-labeled products consistently resolved discrete bands for each chain length—confirming conversion efficiency and substrate purity (Fig. 1b,c). Notably, the polyubiquitin reaction yield and Cy3 labeling efficiency approached ∼100% and ∼98%, respectively, ensuring near-quantitative tracking in downstream single molecule fluorescence assays.

**Fig. 1.**
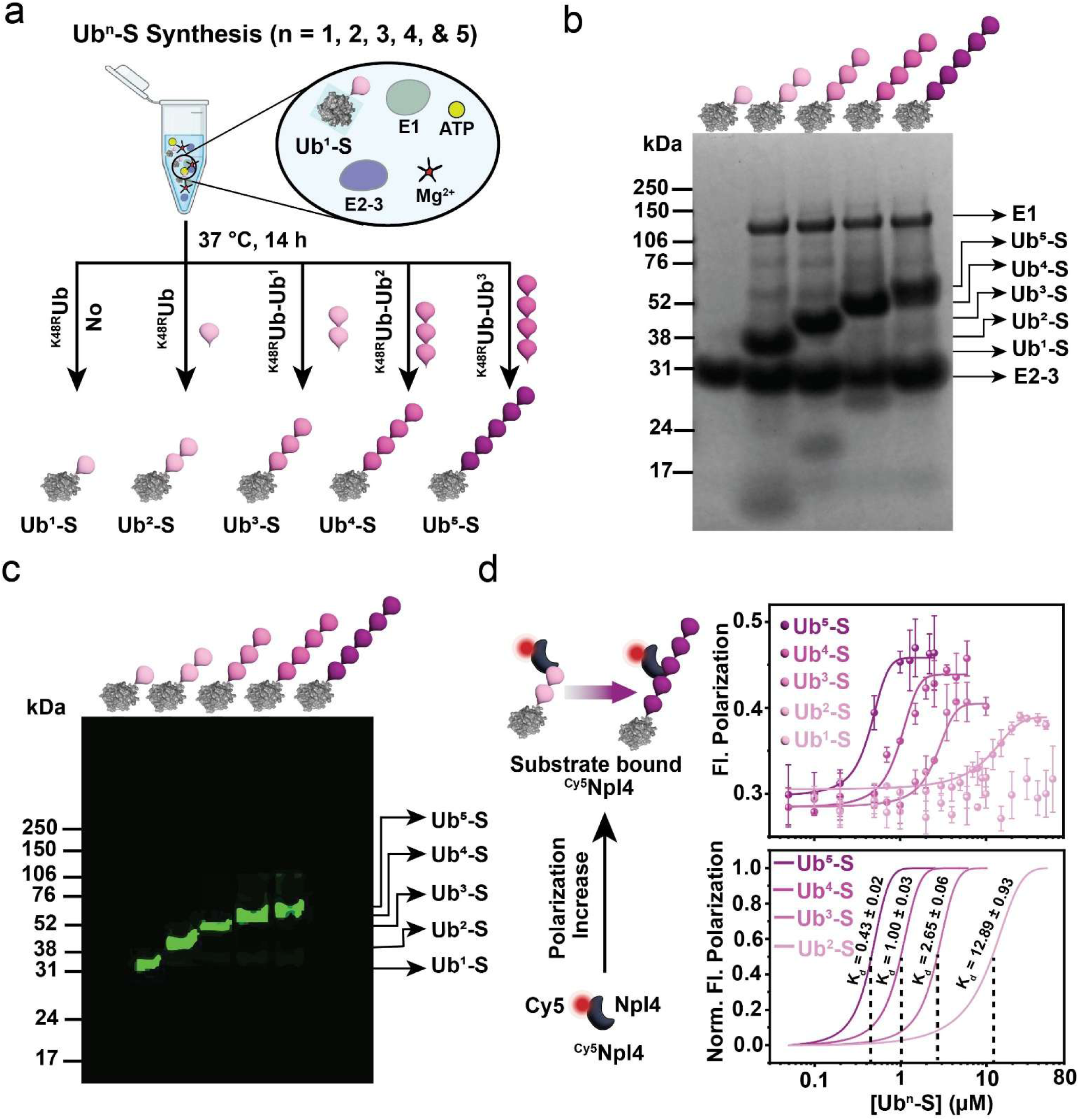
Synthesis and Characterization of Defined-length K48-linked Polyubiquitinated Substrates and Quantification of Npl4–polyubiquitin Interactions. **a** Schematic illustrating the strategy for generating polyubiquitinated substrates with one to five ubiquitin moieties (Ub^1^–S, Ub^2^–S, Ub^3^–S, Ub^4^–S, and Ub^5^–S). Polyubiquitination reactions were carried out in the presence of E1, E2-3 (gp78RING-Ube2g2 chimera), and ATP. b–c SDS–PAGE analysis of purified, biotinylated, and Cy3-labeled polyubiquitin substrates of defined lengths, visualized by Coomassie Blue staining (**b**) and Cy3 fluorescence scanning (**c**), with protein components indicated. **d** Binding of Npl4 to polyubiquitin substrates of varying lengths was quantified by fluorescence polarization using ^Cy5^Npl4 (50 nM; excitation at 640 nm, emission collected between 655–800 nm) titrated with increasing concentrations of Ub^1^–S, Ub^2^–S, Ub^3^–S, Ub^4^–S, and Ub^5^–S, as indicated. Binding curves are presented as raw (top) and normalized (bottom) data. The K_d_ for Ub^1^–S was not determined due to minimal affinity between Npl4 and Ub^1^–S. Data represent mean ± s.d. from three independent experimental replicates.

Next, we quantified how defined polyubiquitin chain lengths influence Npl4 binding using a bulk fluorescence polarization assay. Cy5-labelled Npl4 (^Cy5^Npl4) was titrated with increasing concentrations of Ub^1^–S to Ub^5^–S substrate (Fig. 1d). ^Cy5^Npl4 alone showed low polarization, which increased upon substrate binding. Binding to Ub^1^–S was too weak to determine reliably, consistent with insufficient engagement of Npl4’s ubiquitin-binding elements by a single ubiquitin^37^. In contrast, Ub^2^–S and Ub^3^–S bound detectably, with equilibrium dissociation constants (K_d_) of 12.89 ± 0.93 µM and 2.65 ± 0.06 µM, respectively. Notably, Ub^4^–S and Ub^5^–S showed increasingly tight binding (K_d_ = 1.00 ± 0.03 µM and 0.43 ± 0.02 µM). Overall, affinity improved non-linearly with chain length (∼5-fold from Ub^2^–S to Ub^3^-S, ∼13-fold to Ub^4^–S, and ∼30 -fold to Ub^5^–S), implicating a cooperative, multivalent substrate engagement in which Npl4—known to form a two-ubiquitin clamp^30^—accesses additional, stabilizing interaction surfaces on chains at least four ubiquitins in length.

### smUbiRAD uncovers dynamic multivalent polyubiquitin recognition modes by Npl4

To dissect the mechanism underlying the non-linear increase in affinity between Npl4 and polyubiquitin, we asked whether longer polyubiquitin chains primarily alter encounter frequency (association), residence time (dissociation), or enable additional binding modes. We therefore developed a smUbiRAD to visualize Npl4-polyubiquitin interactions in real time. In this assay, biotinylated Cy3-labelled substrates bearing defined ubiquitin chains (Ub^1^–S to Ub^5^–S) are immobilized on passivated cover glass *via* streptavidin, mapped by TIRF microscopy, and monitored in real time by dual-color imaging to record Cy3 (substrate) and Cy5 (Npl4) signals during repeated binding/dissociation cycles (Fig. 2a). Photobleaching controls verified the presence of single substrate molecules and confirmed that successive single ^Cy5^Npl4 appearances reflect genuine association–dissociation cycles on the same tethered molecule (Fig. 2b).

**Fig. 2.**
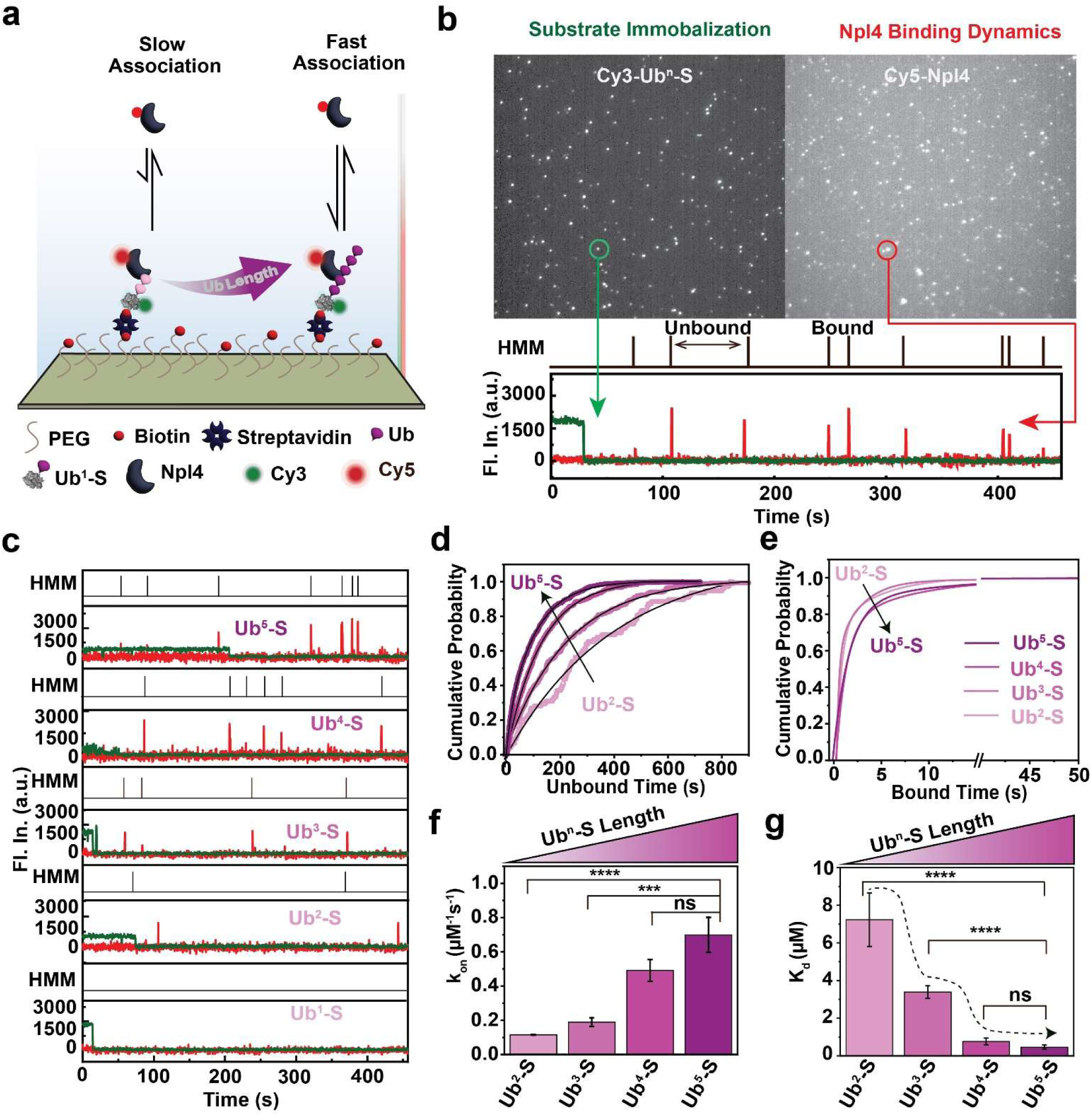
Single-molecule analysis reveals ubiquitin chain length-dependent modulation of Npl4–polyubiquitin interaction dynamics. **a** Schematic of the single-molecule fluorescence assay used to assess Npl4–polyubiquitin interactions. Cy3-labeled, K48-linked polyubiquitin substrates of varying lengths were immobilized on a glass coverslip to determine association and dissociation kinetics for ^Cy5^Npl4 via single molecule total internal reflection fluorescence (smTIRF) microscopy. **b** Representative smTIRF image from the experiment described in (**a**). Left: immobilized polyubiquitin substrate channel (532 nm excitation). Right: ^Cy5^Npl4 binding channel (650 nm excitation). Bottom: representative single-molecule intensity traces. Green traces show single-step photobleaching events for Ub^n^-S substrates (n = 1–5), confirming successful immobilization of individual molecules. Red traces illustrate repetitive binding and unbinding events of ^Cy5^Npl4 on the same substrate. Binding events were analyzed using a two-state Hidden Markov Model (HMM). **c** Representative binding traces of ^Cy5^Npl4 with K48-linked polyubiquitin substrates of different lengths; HMM modeling indicated above each trace. **d-e** Cumulative distributions of unbound (**d**) and bound (**e**) dwell times for ^Cy5^Npl4 interacting with Ub^n^-S substrates. For Ub^2^-S and Ub^3^-S, the distributions were best fit by a single exponential decay model, while Ub^4^-S and Ub^5^-S required a bi-exponential fit. The bi-exponential behavior for longer chains (Ub^4^-S, Ub^5^-S) suggests two distinct Npl4 binding modes, corresponding to interactions with both proximal and distal ubiquitin units. **f-g** Quantification of k_on_ (**f**) and K_d_ (**g**) for ^Cy5^Npl4 binding as a function of polyubiquitin chain length. Statistical significance was determined using a two-tailed Student’s t-test (ns, not significant; 0.05 < p ≤ 0.5; *0.01 < p ≤ 0.05; **0.001 < p ≤ 0.01; ***0.0001 < p ≤ 0.001; ****p ≤ 0.0001).

Single-molecule trajectories revealed a strong dependence of the Npl4 binding kinetics on chain length (Fig. 2c). For Ub^2^–S and Ub^3^–S, unbound and bound dwell-time distributions were well described by single exponentials (Fig. 2d,e), indicating a single dominant interaction mode. Hidden Markov Modelling yielded association rate constant (k_on_) values of 0.115 ± 0.001 μM⁻¹ s⁻¹ (Ub^2^–S) and 0.189 ± 0.025 μM⁻¹ s⁻¹ (Ub^3^–S), with corresponding dissociation rate constant (k_off_) values of 0.827 ± 0.129 s⁻¹ and 0.646 ± 0.142 s⁻¹. The resulting K_d_ (estimating from, k_off_/k_on_) are 7.23 ± 1.42 μM and 3.39 ± 0.33 μM, comparable to our ensemble measurements. In contrast, Ub^4^–S and Ub^5^–S displayed a substantially increasing k_on_ of 0.491 ± 0.064 μM⁻¹ s⁻¹ (Ub^4^–S) and 0.669 ± 0.102 μM⁻¹ s⁻¹ (Ub^5^–S), approaching saturation between four and five ubiquitins (Fig. 2d,f). In parallel, dissociation slowed modestly with length (k_off_ = 0.364 ± 0.042 s⁻¹ for Ub^4^–S; 0.299 ± 0.036 s⁻¹ for Ub^5^–S; Fig. 2e), yielding K_d_ values of 0.763 ± 0.177 μM and 0.462 ± 0.112 μM (Fig. 2g). Notably, Ub^4^–S and Ub^5^–S exhibited two bound-state populations among associating single Npl4 molecules (Fig. 2d,e): their bound dwell-time distributions required bi-exponential fits, revealing a predominant short-lived component (85% of events; ∼1–2 s) and a smaller but reproducible long-lived component (15% of events; ∼8–10 s) (Fig. 2e). This long-lived state emerged only at chain lengths of four or more ubiquitins, identifying a threshold at which Npl4 accesses an additional stabilized binding mode. Given the multiple ubiquitin-interaction elements within Npl4^30^, the appearance of both increasing encounter frequency and stabilized binding is consistent with a multivalent substrate engagement that is sterically or geometrically inaccessible on shorter polyubiquitin chains and becomes feasible only when longer chains can bridge multiple interaction surfaces within the same complex.

Together, our data show that the bulk affinity gain reflects faster capture, slower release, and—critically—a chain-length–licensed long-lived binding mode. More specifically, smUbiRAD demonstrates that full-length human Npl4 decodes K48 linked chain length *via* a kinetic switch at ∼4 ubiquitins, transitioning from predominantly transient binding on short chains to mixed-mode behavior that includes a stabilized long-lived state on Ub^4^–S and Ub^5^–S wherein single Npl4 molecules access an expanded binding surface.

### Ufd1 and p97 specifically enhance substrate retention by Npl4 through hierarchical stabilization

Npl4 and Ufd1 form a stable heterodimer in cells that subsequently assembles with p97 to generate a multivalent ternary complex^25, 26, 29, 38, 39^. Having established that tetra- and penta-ubiquitin chains support high-affinity, multivalent engagement by Npl4, we next tested how adding Ufd1 and p97 reshapes Npl4–substrate dynamics. Using smUbiRAD, we monitored ^Cy5^Npl4 binding to immobilized Ub^5^–S under three conditions: Npl4 alone, the pre-incubated Npl4**–**Ufd1 heterodimer, and the reconstituted Npl4**–**Ufd1**–**p97^WT^ complex (Fig. 3a; Supplementary Fig. 2, Fig. 3; and Supplementary Table 1). Representative single-molecule trajectories displayed repeated ^Cy5^Npl4 binding events in all cases, but residence times increased progressively as components were added (Fig. 3b-d, left). Consistently, intensity distributions shifted toward longer binding events: the long-residence fraction rose from ∼15% for Npl4 alone to ∼40% with Ufd1 and to ∼50% for the full Npl4**–**Ufd1**–**p97^WT^ complex (Fig. 3b-d, right). Thus, cofactor assembly preserves dynamic binding while biasing the system toward the long-lived, multivalent mode.

**Fig. 3.**
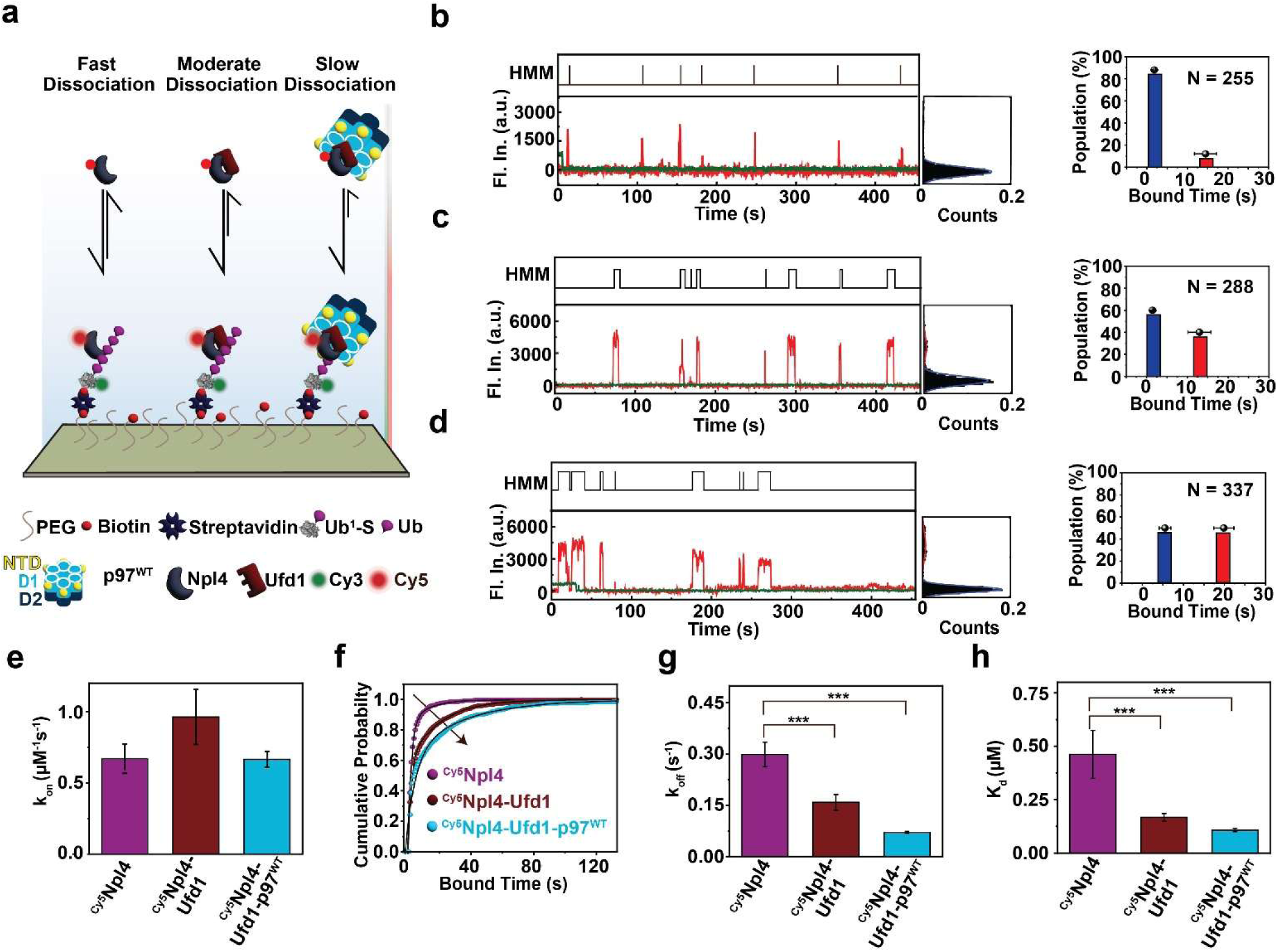
Modulation of Npl4 binding dynamics by Ufd1 and p97 on polyubiquitinated substrates. **a** Schematic of the experimental design for quantifying association and dissociation kinetics of ^Cy5^Npl4 binding to Ub^5^-S substrates under three conditions: Npl4 alone, after assembly with Ufd1, or after assembly with the Ufd1–p97^WT^ complex. **b-d** Representative single-molecule fluorescence intensity trajectories over time, corresponding intensity histogram distribution (left), and the relative amplitude of the long and short binding events (right) of ^Cy5^Npl4 with Ub^5^-S for (**b**) Npl4 alone; (**c**) Npl4 after assembly with Ufd1; and (**d**) Npl4 after assembly with the Ufd1–p97^WT^ complex. HMM analysis is indicated above each trace. Short-lived (blue) and long-lived (red) bound states and their amplitudes are indicated; the fraction of long-lived events increases from ∼15% for Npl4 alone to ∼40% with Ufd1 and ∼50% with Ufd1–p97^WT^. Thus, cofactor assembly preserves dynamic binding while biasing the system toward the long-lived, multivalent mode **e** k_on_ of ^Cy5^Npl4 binding to Ub^5^-S measured under the three conditions described above. **f** Cumulative distributions of bound dwell times of ^Cy5^Npl4 on Ub^5^-S substrate, in the absence and presence of Ufd1 or the Ufd1–p97^WT^ complex. **g-h** Quantification of (**g**) k_off_ and (**h**) K_d_ for ^Cy5^Npl4 binding to Ub^5^-S, in the absence and presence of Ufd1 and Ufd1–p97^WT^, as indicated. Statistical significance was determined using a two-tailed Student’s t-test (ns, not significant; 0.05 < p ≤ 0.5; *0.01 < p ≤ 0.05; **0.001 < p ≤ 0.01; ***0.0001 < p ≤ 0.001; ****p ≤ 0.0001).

Kinetic parameters underscore this ordered stabilization. The k_on_ of ^Cy5^Npl4 with Ub^5^–S was largely unchanged across assemblies, indicating that Ufd1 and p97 do not substantially affect initial substrate capture (Fig. 3e-g; Supplementary Fig. 4). In contrast, dissociation slowed in a stepwise manner: k_off_ decreased from 0.299 ± 0.036 s⁻¹ (Npl4 alone) to 0.159 ± 0.023 s⁻¹ (with Ufd1) and to 0.0716 ± 0.003 s⁻¹ (with both Ufd1 and p97^WT^), yielding corresponding K_d_ values of 0.462 ± 0.112 μM, 0.167 ± 0.018 μM and 0.108 ± 0.008 μM (Fig. 3f-h), respectively. Complementary measurements with ^Cy5^Ufd1 and ^Cy5^p97^WT^ showed weak binding to Ub^5^–S in isolation that strengthened upon complex formation, with the largest reduction in k_off_ observed for the complete assembly (Supplementary Fig. 5, and 6).

Collectively, these findings support a hierarchical model in which Npl4 provides chain-length-sensitive recognition of K48 polyubiquitin, whereas Ufd1 and p97 primarily function as retention factors that extend complex lifetime by suppressing dissociation rather than enhancing encounter^40^.

### Ub^5^–S potently stimulates p97 ATPase

To test whether a defined K48 polyubiquitin signal is sufficient to stimulate p97, we measured ATP activity of p97^WT^ in the context of the Npl4**–**Ufd1 cofactor complex with or without the Ub^5^–S substrate, using Ub^1^–S as a short-chain control (Supplementary Fig. 7a). Ub^1^–S produced little to no change in basal p97 activity, and neither did Npl4**–**Ufd1 alone. In contrast, addition of Ub^5^–S increased ATPase activity by ∼3-fold, comparable to stimulation reported for longer polyubiquitin-conjugated model substrates^21, 29^. These findings indicate that a five-unit K48 chain provides a potent activation cue and that chain architecture—rather than the identity of the attached cargo—is the primary determinant of p97 stimulation through the Npl4–Ufd1 pathway.

We next separated the contributions of the two ATPase rings using Walker A mutants that selectively disrupt nucleotide binding in D1 (K251A) or D2 (K524A) (Supplementary Fig. 3, 7b). The D1-impaired variant retained basal activity but showed only weak Ub^5^–S-dependent stimulation (∼1.6-fold), indicating that D1 is required to reach maximal substrate-driven rate constants. In contrast, disabling D2 exhibited significantly reduced ATP hydrolysis and could not be rescued by cofactors or substrate, establishing D2 as the dominant domain that supports high ATP turnover during activation, and consistent with it forming the major force-generating ring in p97^41^. Consistent with their gain-of-function phenotypes, the MSP-linked variants R155H and A232E—both residing at the NTD-D1 interface and affecting the NTD conformation^19, 35, 42^—displayed ∼3-fold elevated basal ATPase activity and were further stimulated upon cofactor and substrate addition, suggesting a heightened D1 responsiveness to polyubiquitin-triggered activation^43^.

### ATP hydrolysis drives Npl4 cycling in the Npl4–Ufd1–p97 complex

We next asked whether nucleotide processing regulates substrate recognition and cofactor dynamics within the p97 machinery. Prior structural and biophysical work has established that the NTDs of p97 switch between “DOWN” conformations in apo/ADP states and “UP” conformations upon ATP binding, with D1 nucleotide occupancy allosterically controlling NTD positioning (Fig. 4a)^31, 44^. Together with cryo-EM structures showing multiple “seesaw” conformations of Npl4 atop the D1 ring, these observations suggest that ATP binding and/or hydrolysis remodel cofactor engagement by modulating NTD dynamics^28^. We hypothesized that, additionally, nucleotide state may govern Npl4 residence time on polyubiquitin within the assembled Npl4–Ufd1–substrate complex.

**Fig. 4.**
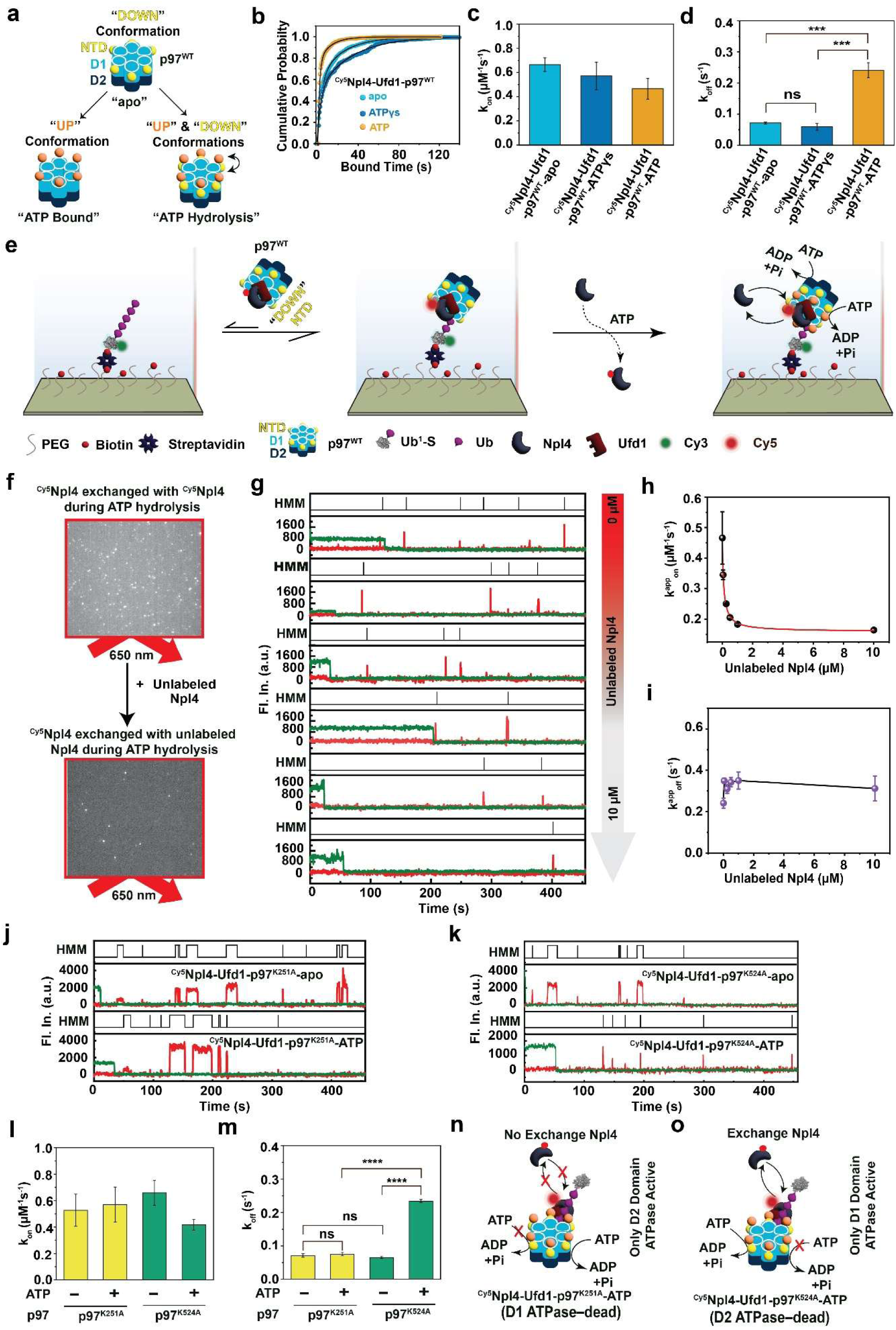
Dynamic Npl4 exchange within the Npl4–Ufd1–p97–Ub^5^-S complex is regulated by conformational changes in p97 during ATP hydrolysis. **a** Schematic illustration of p97^WT^ NTD conformational states (“UP” and “DOWN”) under apo, ATP-bound, and ATP hydrolysis conditions. **b** Cumulative distributions of bound dwell times of ^Cy5^Npl4 (after assembly with Npl4–Ufd1–p97^WT^ complexes) on the Ub^5^-S substrate under apo (light blue), ATPγS (dark blue), and ATP (gold) conditions, highlighting altered Npl4 stability depending on the nucleotide state. **c** k_on_ and **d** k_off_ of ^Cy5^Npl4 (after assembly with Npl4–Ufd1–p97^WT^ complexes) on the Ub^5^-S substrate under apo, ATPγS, and ATP conditions, demonstrating nucleotide-dependent exchange kinetics. **e** Schematic of ^Cy5^Npl4 exchange by unlabeled Npl4 within the Npl4**–**Ufd1**–**p97^WT^**–**Ub^5^-S complex during ATP hydrolysis. **f** Representative smTIRF image showing ^Cy5^Npl4 (top, [unlabeled Npl4] = 0 μM) and in the presence of unlabeled Npl4 (bottom, [unlabeled Npl4] = 10 μM) upon excitation at 650 nm. **g** Representative single-molecule traces depicting ^Cy5^Npl4 binding and exchange events (red) as increasing concentrations of unlabeled Npl4 are introduced, with HMM analysis displayed above each trace (black). Green traces show single-step photobleaching of the Cy3-labeled substrate, confirming a single substrate molecule for each spot in the green channel. **h-i** Apparent association rate constant k^app^_on_ (**h**) and apparent dissociation rate constant, k^app^_off_ (**i**) for ^Cy5^Npl4 in the Npl4–Ufd1–p97^WT^ complex on the Ub^5^-S substrate at different concentrations of unlabeled Npl4. **j-k** Representative single-molecule time trajectories of ^Cy5^Npl4 binding to Ub^5^-S substrate in the Npl4–Ufd1–p97 complexes containing p97^K251A^ (**j**) and p97^K524A^ (**k**) in the absence (top) and presence (bottom) of ATP; HMM analysis is indicated above each trace. **l-m** The k_on_ (**l**) and k_off_ (**m**) for ^Cy5^Npl4 binding to Ub^5^-S substrate in Npl4–Ufd1–p97 complexes congaing p97^K251A^ or p97^K524A^, in the absence and presence of ATP, indicating domain-specific control of Npl4 dynamics. **n** Schematic showing lack of Npl4 exchange in the Npl4–Ufd1–p97^K251A^–Ub^5^-S complex during ATP hydrolysis, specifically at the D2 domain. **o** Schematic showing Npl4 exchange in the Npl4–Ufd1–p97^K524A^–Ub^5^-S complex during ATP hydrolysis, specifically at the D1 domain. Statistical significance was determined using a two-tailed Student’s t-test (ns, not significant; 0.05 < p ≤ 0.5; *0.01 < p ≤ 0.05; **0.001 < p ≤ 0.01; ***0.0001 < p ≤ 0.001; ****p ≤ 0.0001).

To test this hypothesis, we employed smUbiRAD to monitor ^Cy5^Npl4 in preassembled Npl4–Ufd1–p97^WT^ complexes bound to immobilized Ub^5^–S under three conditions: nucleotide-free (apo), ATPγS (non-hydrolysable analogue; ATP-bound state), and ATP (hydrolysis enabled)^45^. The k_on_ of ^Cy5^Npl4 was largely insensitive to nucleotide state (Fig. 4b; Supplementary Fig. 8), indicating that initial substrate capture is not strongly nucleotide dependent. In contrast, dissociation accelerated specifically during ATP hydrolysis: k_off_ increased ∼4-fold in ATP conditions relative to apo or ATPγS, in which Npl4 remained more stably associated (Fig. 4c,d; Supplementary Fig. 8b,c). Thus, ATP hydrolysis promotes Npl4 turnover within the complex. We then asked whether this hydrolysis-dependent destabilization reflects disassembly of the entire assembly or selective exchange of Npl4. Single-molecule measurements showed that Ufd1 and p97 remained stably bound to Ub^5^–S during ATP turnover and did not display a comparable increase in dissociation (Supplementary Fig. 5, and 6). These results support a model in which ATP hydrolysis selectively triggers Npl4 turnover, whereas its heterodimer partner Ufd1 remains bound and the Ufd1–p97 complex persists as a substrate-bound scaffold.

We next tested this exchange model using a competition assay. Increasing concentrations of unlabeled Npl4 were added to reactions containing ^Cy5^Npl4, Ub^5^–S, Ufd1 and p97^WT^ under ATP-hydrolysis conditions (Fig. 4e). Unlabeled Npl4 reduced ^Cy5^Npl4 rebinding frequency and lengthened unbound intervals in a concentration-dependent manner (Fig. 4f,g; Supplementary Fig. 9). Consistent with competitive replacement^46^, the apparent association rate constant (k_on_^app^) for ^Cy5^Npl4 decreased sharply as the unlabeled Npl4 increased, accompanied by a modest rise in the apparent dissociation rate constant (k ^app^) (Fig. 4f,g; Supplementary Fig. 9).

Together, these data show that ATP hydrolysis drives cyclical recruitment and release of Npl4 while preserving the integrity of the Ufd1–p97 scaffold on the polyubiquitin substrate.

### D1-ATP hydrolysis powers Npl4 cycling

The regulatory D1 and power-stroke D2 ATPase domains both contribute to the overall ATPase activity of p97^47, 48^. Our observation that ATP turnover accelerates Npl4 release from the Npl4–Ufd1–p97–Ub^5^–S complex prompted us to ask which ATPase ring (D1 or D2) supplies the signal that gates Npl4 dynamic exchange. To dissect ring-specific contributions, we used Walker A mutants—p97^K251A^ and p97^K524A^—to selectively impair ATP binding and hydrolysis in D1 and D2, respectively (Supplementary Fig. 3, and 7). In the p97^K524A^ mutant, which impairs D2 while preserving D1 function, ATP addition increased the Npl4 k_off_ by ∼3.6-fold, from 0.065 ± 0.003 s⁻¹ to 0.234 ± 0.005 s⁻¹, closely resembling the behavior of p97^WT^under ATP-hydrolysis conditions (Fig. 4j-m, and Supplementary Fig. 10). In contrast, in the p97^K251A^ mutant—defective in D1 ATP processing—the Npl4–Ufd1–p97^K251A^–Ub^5^–S complex showed no significant change in Npl4 binding kinetics in the absence (apo) and presence of ATP, with a k_off_ of 0.071 ± 0.005 s⁻¹ (apo) and 0.075 ± 0.005 s⁻¹ (ATP); as before, the k_on_ also remained unchanged (Fig. 4j-m, and Supplementary Fig. 10). These results demonstrate that ATP hydrolysis in the D1 ring—but not the D2 ring—is both necessary and sufficient to drive Npl4 exchange within the Npl4–Ufd1–p97–Ub^5^–S complex (Fig. 4n-o). Furthermore, given the mean Npl4 residence time of ∼4 s during ATP turnover, our data are consistent with a model in which coordinated D1 chemo-mechanical transitions are temporally coupled to discrete Npl4 turnover events, establishing D1 as a dedicated cofactor-exchange engine and Npl4 cycling as a gating factor for iterative coupling to downstream D2-driven substrate processing^31^.

### MSP-linked p97 mutants promote enhanced substrate engagement and hyperactivity

Notably, numerous pathogenic p97 mutations cluster at the NTD–D1 interface and are proposed to bias the NTD toward an “UP” conformation, yielding gain-of-function activity^19, 35, 49^. With D1 ATP hydrolysis identified as the trigger for Npl4 exchange, we asked how MSP variants reshape this cycle. Using smUbiRAD, we compared p97^WT^ with the mutants R155H and A232E in complexes with Npl4–Ufd1 on Ub^5^–S across apo, ATPγS-bound and ATP-hydrolysis conditions (Fig. 5a; Supplementary Fig. 11).

**Fig. 5.**
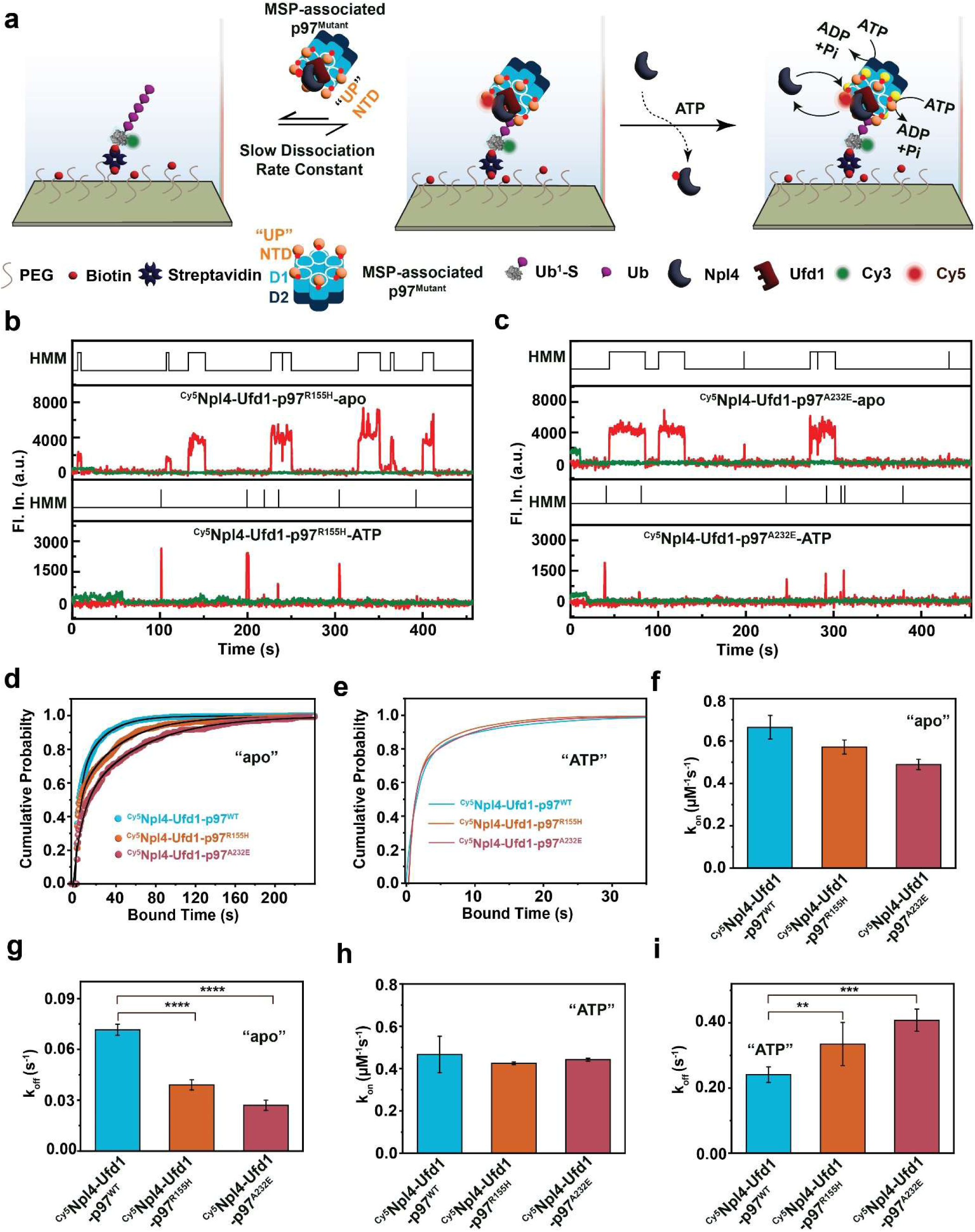
MSP-associated p97 mutants R155H and A232E increase substrate affinity and Npl4–Ufd1–p97 complex dynamics. **a** Schematic of MSP-associated p97 mutants R155H and A232E in complex with cofactors Npl4–Ufd1 and the Ub^5^-S substrate in the “apo” or “ATP” states. In the “apo” state, p97 mutants adopt an N-terminal domain (NTD) “UP” conformation, whereas p97^WT^ assumes an NTD “DOWN” conformation. **b-c** Representative single-molecule fluorescence traces showing ^Cy5^Npl4 (red) binding to Ub^5^-S after assembly with (**b**) Npl4–Ufd1–p97^R155H^ or (**c**) Npl4–Ufd1–p97^A232E^. Upper and lower panels correspond to the “apo” and “ATP” states, respectively. HMM state assignments (black) are indicated above each trace. Green traces show single-step photobleaching of the Cy3-labeled substrate, confirming a single substrate molecule for each spot in the green channel. **d** Cumulative distributions of ^Cy5^Npl4 bound dwell times (after assembly with Npl4–Ufd1–p97) on Ub^5^-S in the absence of ATP for p97^WT^, p97^R155H^, or p97^A232E^, highlighting altered complex stability of the mutants relative to p97^WT^. **e-f** Comparison of kinetic parameters (association rate constant, k_on_ and dissociation rate constant, k_off_) for ^Cy5^Npl4 binding to Ub^5^-S after assembly with Npl4–Ufd1–p97 in the absence of ATP: (**e**) k_on_ and (**f**) k_off_ for p97^WT^ (cyan), p97^R155H^ (orange), and p97^A232E^ (red). **g** Cumulative distributions of ^Cy5^Npl4 bound dwell times (after assembly with Npl4–Ufd1–p97) on Ub^5^-S in the presence of ATP for p97^WT^, p97^R155H^, and p97^A232E^. **h-i** Comparison of kinetic parameters for ^Cy5^Npl4 binding to Ub^5^-S after assembly with Npl4–Ufd1–p97 in the presence of ATP: (**h**) k_on_ and (**i**) k_off_ for p97^WT^, p97^R155H^, or p97^A232E^. Statistical significance was determined using a two-tailed Student’s t-test (ns, not significant; 0.05 < p ≤ 0.5; *0.01 < p ≤ 0.05; **0.001 < p ≤ 0.01; ***0.0001 < p ≤ 0.001; ****p ≤ 0.0001).

In the nucleotide-free state, both mutant complexes retained Npl4 more strongly than WT, exhibiting slower dissociation (k_off_ = 0.039 ± 0.003 s⁻¹ for R155H and 0.027 ± 0.003 s⁻¹ for A232E versus 0.0716 ± 0.003 s⁻¹ for WT) (Fig. 5b-g). Consistent with this, their equilibrium affinities were higher (K_d_ = 0.068 ± 0.002 μM and 0.056 ± 0.007 μM) than WT (0.108 ± 0.008 μM) (Supplementary Fig. 11), matching structural models in which these variants favor an NTD “UP”-like state that strengthens cofactor engagement and substrate recognition^19, 49^.

Upon ATPγS binding, dissociation and affinity values converged across WT and mutants, indicating that ATP binding produces a similar Npl4-bound state (Supplementary Fig. 8). In contrast, during ATP hydrolysis, Npl4 residence times decreased for both mutants relative to WT (mean dwell times ∼3.10 s for R155H and ∼2.45 s for A232E versus ∼4.2 s for WT), translating into increased dissociation rate constants (k_off_ = 0.34 ± 0.06 s⁻¹ and 0.41 ± 0.034 s⁻¹ versus 0.241 ± 0.024 s⁻¹ for WT; Fig. 5e-i). Association rate constants were largely unchanged, but the faster turnover increased the apparent K_d_ values for the mutants (0.741 ± 0.145 μM and 0.759 ± 0.088 μM) relative to WT (0.529 ± 0.063 μM), consistent with accelerated Npl4 exchange in the complex.

Taken together, across nucleotide states mutant and WT dwell times were similar under apo and ATPγS conditions but specifically shortened upon ATP hydrolysis in the disease variants (Fig. 5; Supplementary Fig. 8, and 11). These results indicate dysregulated cofactor cycling as a kinetic consequence of MSP mutations: they exhibit stronger substrate engagement at rest (apo) coupled to faster D1-driven Npl4 turnover during active hydrolysis, consistent with a gain-of-function mechanism that may lead to weaker D1-D2 coupling and aberrant substrate processing, and resulting in the observed proteostasis imbalance^31^.

## Discussion

This study offers a mechanistic advance in our understanding of p97-mediated proteostasis by leveraging single-molecule measurements to resolve the choreography of cofactor engagement, substrate recognition and ATPase-driven dynamics. Using smUbiRAD, we directly observed polyubiquitin decoding and cofactor behavior in real time and find that the human p97 system is more modular and kinetically tuned than previously appreciated^13, 23, 30, 50^, with Npl4 acting as the primary chain-length sensor, Ufd1 and p97 as substrate retention factors, and D1 ATPase activity as a dedicated cofactor-exchange engine that enforces Npl4 cycling to regulate D2 power-stroke progression (Fig. 6).

**Fig. 6.**
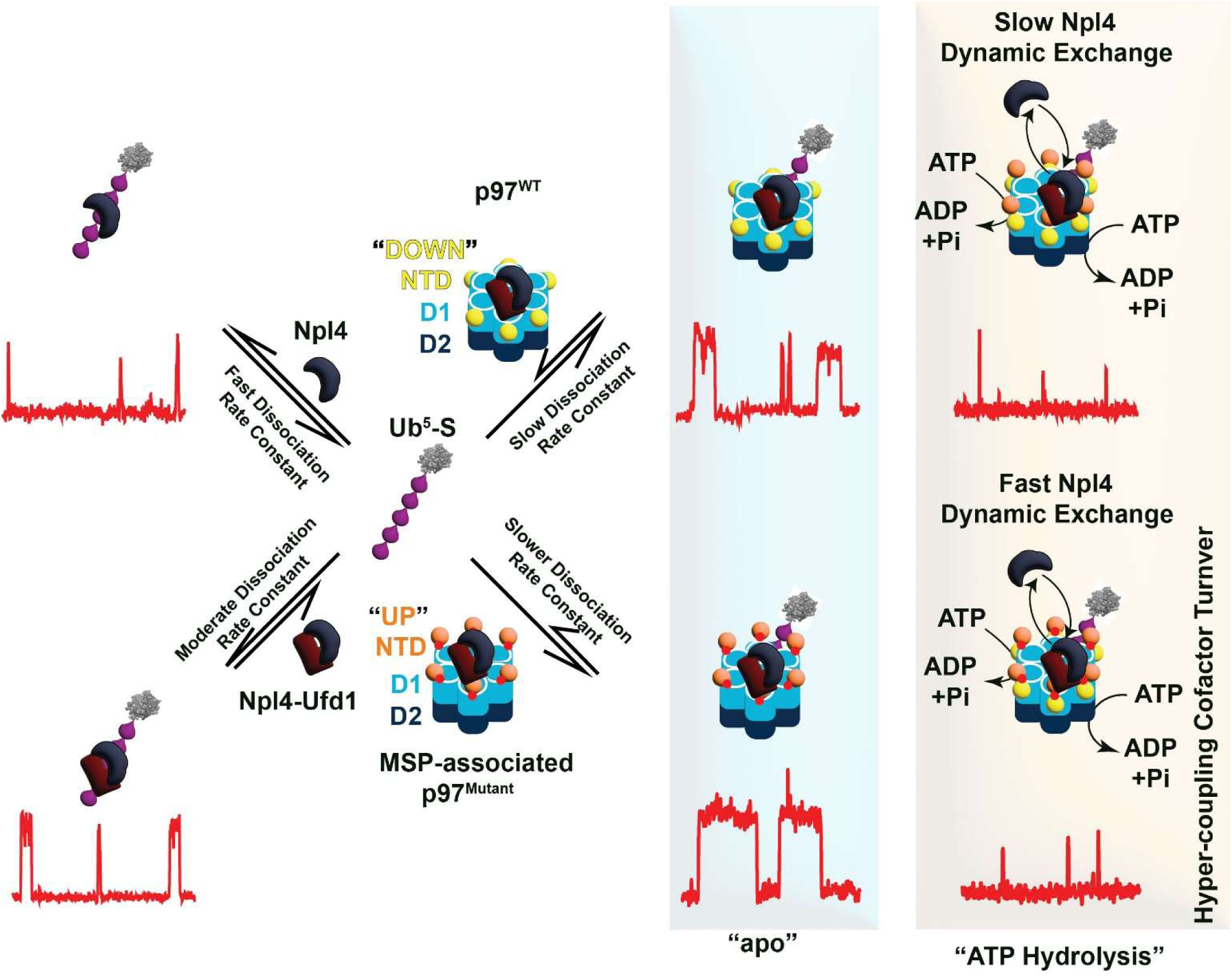
Integrative model of chain-length decoding, D1-gated cofactor cycling and hyperactivation of Npl4 exchange associated with MSP-linked p97 variants. smUbiRAD measurements support a kinetic model in which Npl4 acts as the primary reader of K48-linked polyubiquitin. Assembly with Ufd1 and p97 stabilizes substrate-bound complexes mainly by suppressing Npl4 dissociation, thereby increasing residence time without markedly changing association rate constants. Upon ATP turnover, p97 selectively promotes Npl4 exchange while Ufd1 and p97 remain associated with the substrate, forming a persistent scaffold. Domain-selective perturbations indicate that D1 nucleotide processing is necessary and sufficient to gate Npl4 turnover. MSP-linked p97 variants (R155H and A232E) tighten cofactor engagement in the apo state yet accelerate ATP-driven Npl4 exchange, consistent with hyperactive cofactor cycling as a mechanism of pathogenic dysregulation.

One of our prominent observations is that human Npl4 exhibits a threshold-like response to K48 chain length. Bulk measurements show a strong, non-linear increase in affinity as chains extend beyond three ubiquitins. smUbiRAD reveals the kinetic basis of this behavior and identifies a qualitative transition at ≥ Ub^4^, where Npl4 binding becomes heterogeneous and a distinct, long-lived population emerges (Fig. 2)^24^. Specifically, Npl4 shifts from transient, largely single-site interactions on short chains to stabilized binding on tetra- and penta-ubiquitin, consistent with an intrinsic “counting” mechanism for discriminating degradation signals. This non-linear affinity gain and long-lived state imply cooperative engagement of multiple ubiquitin-binding motifs within Npl4, highlighting chain architecture—not simply length or cargo identity—as a key determinant for p97 targeting^24, 30^. The fact that still only a single Npl4 molecule binds to the elongated ubiquitin chain suggests that avidity is facilitated by sliding or short-range hopping between concatenated binding sites.

Cofactor assembly is expected to further refine this recognition landscape^14, 30^. Although initial association rate constants remain largely unchanged, incorporation of Ufd1 and p97 progressively suppresses dissociation, producing hierarchical stabilization (Fig. 3). This kinetic partitioning positions Npl4 as the primary ubiquitin reader, whereas Ufd1 and p97 mainly extend bound-state lifetime, committing an initial recognition event into a long-lived processing platform. Such modular stabilization provides a mechanism to tune substrate dwell time without altering chain-length specificity and is likely critical for coupling recognition to unfolding under cellular conditions where multiple substrates compete for unfoldase engagement. Indeed, recent work demonstrated accessory adaptor proteins such as FAF1 interact with Ufd1 do not have a marked impact on substrate recognition, but rather increase unfolding activity and downstream proteolysis^24, 27^.

A key conceptual advance from our real-time measurements is that ATP hydrolysis in the D1 ring, but not D2, is both necessary and sufficient to trigger rapid Npl4 exchange while Ufd1 and p97 remain scaffolded on the substrate (Fig. 4). This links D1 chemo-mechanical cycling to cofactor turnover and provides a mechanistic basis for the nucleotide-driven NTD “UP/DOWN” switching observed structurally^31, 51^. The finding that D2 ATPase activity is dispensable for Npl4 exchange but required for robust ATPase stimulation and substrate processing supports a division of labor in which the D1 ring orchestrates timing and cofactor flux, whereas the D2 ring supplies high-power mechanical work^52^. Npl4 exchange may thus function as a reset mechanism that prevents kinetic trapping and enables iterative substrate engagement during processing, endowing the Npl4–Ufd1–p97 machinery with an escapement-like reciprocal gating mechanism that permits progression only after completion of the prior unfolding cycle, reminiscent of mechanisms utilized by the proteasome and other AAA+ ATPase machines^53, 54^.

Multisystem proteinopathy-associated p97 mutations mapped to the NTD–D1 interface (R155H, and A232E) further underscore this control logic. These variants tighten Npl4 association in the resting state, yet accelerate Npl4 turnover during ATP hydrolysis (Fig. 5), generating a hyper-exchange regime superimposed on elevated ATPase activity that dampens D1–D2 ring coupling. Our findings thus suggest a kinetic basis for aberrant hyperactive substrate processing in multisystem proteinopathy^16, 55^.

In summary, our findings redefine the Npl4–Ufd1–p97 machinery as a dynamic, chain-length-gated decoding module in which recognition, retention and turnover are distributed across cofactors and ATPase rings (Fig. 6). Npl4 emerges as a central component involved in both proper substrate selection by human VCP/p97 and gating the coupling of p97’s D1–D2 ATPase rings through D1-enforced exchange. Furthermore, smUbiRAD provides a generalizable framework to probe how other ubiquitin architectures, cofactors and modulators reshape this kinetic landscape. Future work extending this methodology to branched or mixed-linkage chains, diverse client proteins, and additional disease alleles should clarify how ubiquitin code complexity and cofactor misregulation converge to result in neurodegeneration and cancer, potentially guiding the design of therapies that normalize cofactor cycling rather than fully suppressing p97 function under conditions of aberrant protein quality control.

## Methods

### Plasmid construction

The Ub^1^-SNAP-6xHis plasmid was constructed by introducing ubiquitin N-terminally fused into the pSNAP-CaaX plasmid (NEB #N9180). The 6xHis-TEV-gp78-Ube2g2 plasmid, generously provided by Andreas Martin, employs the pET-28a vector for protein expression in *E. coli*^14^. Coding sequences for 6xHis-TEV-Npl4 and 6xHis-TEV-Ufd1 were obtained from Addgene (#117108 and #117107, respectively) and subcloned into the pET-28a system, consistent with the method used for the 6xHis-TEV-gp78-Ube2g2 construct. Each vector includes an N-terminal TEV protease cleavage site followed by a 6xHis affinity tag. Human p97 (wild-type or mutants) was cloned into a pET-28a bacterial expression vector containing a C-terminal HA tag and a six-histidine tag, separated by a flexible glycine/serine linker (sequence: GGGGSGGGGSGGGGSGGGGSGGGGSG).

### Purification of monoubiquitin substrate (Ub^1^-S)

The linear fusion protein, consisting of ubiquitin linked at its C-terminus to the N-terminus of SNAP-tag (Ub^1^-S), was produced and purified using standard methodologies^21, 56^. *E. coli* BL21 star (DE3) cells were transformed with the appropriate plasmid, cultivated in Terrific Broth (TB) at 37 °C until reaching an optical density at 600 nm (OD_600_) of 1.0, and then induced with 1 mM IPTG, followed by overnight incubation at 18 °C. Cells were collected by centrifugation at 5,000 rcf and resuspended in lysis buffer (50 mM HEPES, pH 7.4; 200 mM KCl; 20 mM imidazole; 10 mM MgCl₂; 5% glycerol; and a protease inhibitor cocktail). The cells were lysed *via* sonication, and the resulting lysate was clarified by centrifuging at 35,000 rcf. The supernatant was incubated with Ni-NTA resin for one hour to enable affinity purification. After extensive washing with lysis buffer, Ub^1^-S was eluted using lysis buffer containing 350 mM imidazole. The eluted protein sample was dialyzed overnight at 4 °C in storage buffer (25 mM HEPES, pH 7.4; 200 mM KCl; 10 mM MgCl₂; 10% glycerol) to remove imidazole. The purified Ub^1^-S protein, now free of imidazole, was divided into aliquots, quickly frozen in liquid nitrogen, and stored at –80 °C until further use.

### Purification of the E2-E3 chimera gp78RING-Ube2g2

The E2-E3 chimera gp78RING-Ube2g2 was produced and purified according to established protocols^14, 21^. In summary, *E. coli* BL21 star (DE3) cells were transformed with the relevant plasmid and cultured in Terrific Broth (TB) at 37 °C until an OD_600_ of 0.6 was reached. Protein expression was induced by adding 0.4 mM IPTG, followed by overnight incubation at 25 °C. The harvested cells were collected by centrifugation and resuspended in lysis buffer (50 mM Tris, pH 7.4; 500 mM KCl; 5 mM MgCl2; 20 mM imidazole; 5% glycerol; 2 mM β-mercaptoethanol; supplemented with protease inhibitors). Cells were disrupted using sonication, and the lysate was clarified *via* centrifugation at 27,000 rcf and filtered through a 0.45 µm membrane. The resulting clarified lysate was incubated with Ni-NTA resin for one hour to ensure binding of the 6xHis-tagged protein, after which the resin was thoroughly washed with lysis buffer. Target proteins were eluted using lysis buffer supplemented with 350 mM imidazole. The eluted protein sample was subjected to TEV protease treatment overnight at 4 °C in storage buffer (20 mM HEPES, pH 7.4; 250 mM KCl; 1 mM MgCl2; 5% glycerol; 0.5 mM TCEP), to remove the N-terminal 6xHis-tag. On the following day, the mixture was applied to fresh Ni-NTA resin to separate the tag-free gp78RING-Ube2g2 chimera from uncleaved protein and TEV protease (here, we used 6xHis TEV-protease). The unbound fraction containing the purified chimera was collected, aliquoted, quickly frozen in liquid nitrogen, and stored at –80 °C for use in further polyubiquitination assays.

### Purification of cofactor proteins Npl4 and Ufd1

For co-expression experiments, Ufd1, bearing a 6xHis tag, and untagged Npl4 were used. Equal amounts of Ufd1 and Npl4 plasmids were co-transformed into *E. coli* cells^28^. When expressing cofactors individually, each plasmid was introduced and expressed separately. In all cases, *E. coli* BL21 star (DE3) cells were transformed with the respective plasmid and cultured in Terrific Broth (TB) at 37 °C until an OD_600_ of 1.0 was reached. Protein expression was induced with 0.4 mM IPTG and the cultures were incubated overnight at 18 °C.

Following expression, *E. coli* cells were collected by centrifugation at 4,000 rcf and resuspended in lysis buffer (50 mM HEPES, pH 7.4; 250 mM KCl; 20 mM imidazole; 1 mM MgCl2; 5% glycerol; 0.5 mM TCEP) supplemented with protease inhibitors. Cells were lysed using sonication, and insoluble material was removed *via* centrifugation at 30,000 rcf, followed by filtration through a 0.45 μm membrane. The clarified lysate was incubated with Ni-NTA resin for one hour to capture His-tagged proteins, before transfer to a chromatography column. After extensive washing with lysis buffer, bound proteins were eluted with lysis buffer containing 350 mM imidazole. The eluted was treated overnight at 4 °C with TEV protease in storage buffer (25 mM HEPES, pH 7.4; 250 mM KCl; 1 mM MgCl2; 5% glycerol; 0.5 mM TCEP) to remove the 6xHis-tag from the C-terminus. A second round of Ni-NTA affinity purification allowed for separation of the cleaved, tag-free proteins from uncleaved material and TEV protease. Flow-through fractions containing untagged Npl4 and Ufd1 were collected. Residual imidazole was removed by buffer exchange using centrifugal concentration and dilution at 2,000 rcf. The purified cofactor proteins were aliquoted, flash-frozen in liquid nitrogen, and stored at –80 °C for subsequent applications.

### Purification of p97^WT^, p97^K251A^, p97^K524A^, p97^R155H^, and p97^A232E^

WT p97 (p97^WT^), engineered Walker A mutants in the D1 (p97^K251A^) and D2 (p97^K524A^) domains, as well as disease-associated variants (p97^R155H^ and p97^A232E^), were each expressed and purified according to standard laboratory protocols^57^. Cultures were incubated at 37 °C with shaking at 250 rpm until an OD_600_ of 0.6 was achieved. Protein production was then induced by adding 0.5 mM IPTG, and cells were further incubated for 14 hours at 25 °C. Following expression, cells were harvested by centrifugation at 5,000 rcf and resuspended in lysis buffer containing 50 mM Tris-HCl, pH 8; 300 mM NaCl; 0.5 mM β-mercaptoethanol; 1 mM ATP; and a protease inhibitor cocktail. The cells were lysed by sonication, and the resulting lysate was clarified by ultracentrifugation at 50,000 rcf to remove cell debris. The supernatant was applied to Ni-NTA agarose resin for affinity purification, followed by washing steps and elution with lysis buffer containing 350 mM imidazole. The eluted protein fraction was dialyzed overnight at 4 °C in storage buffer (20 mM HEPES, pH 7.4; 250 mM KCl; 1 mM MgCl2; 10% glycerol) using a 10 kDa molecular weight cutoff membrane to eliminate residual contaminants. To ensure the removal of imidazole, buffer exchange was performed by repeated concentration and dilution at 5,000 rcf for 30 minutes. The final purified p97 variants were aliquoted, flash-frozen in liquid nitrogen, and stored at – 80 °C for subsequent experimental use.

### Biotinylated and Cy3 labeled with Ub^1^-S

Purified Ub^1^-S was biotinylated using SNAP-Biotin (NEB, Catalog #S9110S), which covalently attaches biotin to the Cys-145 residue on the SNAP-tag, as previously described^58, 59^. Following biotinylation, Ub^1^-S was labeled with a fluorescent Cy3 probe by reacting the protein to the amine-reactive Cy3-NHS ester (Lumiprobe, Catalog #11020), following established protocols (Fig S1). For both reactions, a 10:1 molar excess of SNAP-Biotin and Cy3-NHS ester (prepared in DMSO) relative to Ub^1^-S was used. The labeling proceeded for 1 hour at room temperature in 1× PBS, pH 7.4. To remove excess reagents and purify the Cy3- and biotin-modified Ub^1^-S conjugates, samples were processed using Zeba Spin desalting columns (Thermo Fisher, Catalog #89882, 7K MWCO) according to the manufacturer’s instructions. The degree of Cy3 labeling, quantified by measuring absorbance at 280 nm (for protein, Ub^1^-S) and 550 nm (for Cy3) with a spectrophotometer, consistently exceeded 98%. The doubly modified Ub^1^-S was aliquoted, flash-frozen in liquid nitrogen, and stored at –80 °C until further use.

### Poly-ubiquitination reaction for the preparation of Ub^1^-S, Ub^2^-S, Ub^3^-S, Ub^4^-S, Ub^5^-S

Polyubiquitinated Ub^1^-S derivatives with increasing chain lengths (Ub^2^-S through Ub^5^-S) were synthesized *in vitro* using the following reaction conditions^21^. Cy3-labeled and biotinylated Ub^1^-S fusion protein was incubated at 30 μM with 1 μM E1 enzyme, 20 μM gp78RING-Ube2g2 E2–E3 chimera, and 40 μM K-48-linked modified ubiquitin substrates (such as ^K48R^Ub, ^K48R^Ub-Ub^1^, ^K48R^Ub-Ub^2^, and ^K48R^Ub-Ub^3^, for synthesis of Ub^2^-S, Ub^3^-S, Ub^4^-S, and Ub^5^-S, respectively) in 20 mM HEPES (pH 7.4), supplemented with 10 mM ATP and 10 mM MgCl₂ (Fig. 1a and Supplementary Fig. 1b)^21^. The reaction proceeded at 37 °C for 14 hours, with incremental addition of ubiquitin during the first 8 hours to optimize chain assembly. To isolate polyubiquitinated fusion proteins from unincorporated ubiquitin and other reaction components, the reaction mixture was incubated with Ni-NTA resin; only the substrate protein retained a 6xHis-tag for affinity capture. Bound proteins were eluted with buffer containing 300 mM imidazole and subsequently stored in 20 mM HEPES (pH 7.4), 250 mM KCl, 1 mM MgCl₂, 1 mM tris(2-carboxyethyl) phosphine (TCEP), and 5% glycerol. The various polyubiquitinated products were buffer-exchanged to remove residual imidazole, concentrated using centrifugal filtration units, aliquoted, and flash-frozen in liquid nitrogen for subsequent analyses and stored at -80 °C.

### Cy5 labeling of Npl4, Ufd1, and p97^WT^

Purified Npl4, Ufd1, and p97 proteins were chemically labeled with Cy5-NHS ester dye (Lumiprobe, Cat. #13020) *via* amine-reactive conjugation. For each protein, labeling was performed at a 1:10 molar ratio of protein to dye in a total volume of 50 µL labeling buffer (1× PBS, pH 7.4; Thermo Fisher Scientific, Cat. #10-010-049), resulting in a final protein concentration of approximately 10 µM^60^. The reaction mixture was incubated at room temperature for 1 hour^60^. Unreacted excess dye was removed through two rounds of purification using Zeba™ Spin Desalting Columns (7 kDa MWCO, Thermo Fisher Scientific, Cat. #89882). Labeling efficiency was assessed spectrophotometrically using absorbance readings at ∼280 nm (for protein) and ∼650 nm (for Cy5). Labeling consistently approached 100%, with an average 1:1 molar incorporation of Cy5 per protein molecule. Labeled proteins were then aliquoted, flash-frozen in liquid nitrogen, and stored at −80 °C until use.

### Fluorescence polarization assays

Fluorescence polarization was used to determine the binding affinities (the equilibrium dissociation constant, K_d_) between Cy5-labeled Npl4 (^Cy5^Npl4) and polyubiquitin substrates of varying lengths (Ub^1^-S, Ub^2^-S, Ub^3^-S, Ub^4^-S, and Ub^5^-S) (Fig. 1d). Fluorescence polarization measurements were performed using a Horiba Scientific fluorometer (Fluoromax Plus-C) in a 0.1 cm path length cuvette at a constant temperature of 25 °C. Each binding assay was conducted in triplicate, and experimental data represent the average of three independent scans. For the assay, 30 nM ^Cy5^Npl4 was titrated with increasing concentrations of the respective polyubiquitin substrates, as marked in the figures and main text (Fig. 1d). Fluorescence polarization readings were collected using excitation and emission slit widths of 3 nm for each, with excitation at 640 nm and emission collected from 655 nm to 800 nm. Instrument signal and reference normalization were applied (S1C and R1C enabled), and spectra are presented as the sample-detector signal divided by the reference-detector signal (S1C/R1C). Binding affinities (K_d_) were determined by fitting the data with a sigmoidal model using Origin software.

### ATPase assays

ATPase activity was assessed using a protocol adapted from previously published methods^61, 62^. Assays were conducted in untreated microplates (Greiner BioOne, Cat. #655101) with 40 µL reaction mixtures containing 30 nM p97 hexamer (for all mutant variants), 300 nM Npl4**–**Ufd1 complex, and/or 300 nM substrates (Ub^1^-S and Ub^5^-S) in ATPase assay buffer (25 mM HEPES, pH 7.4, 100 mM KCl, 3 mM MgCl₂, 1 mM TCEP, and 0.1 mg/mL BSA, 0.05% Triton-X). Reactions were preincubated at 37 °C for 10 min, followed by the addition of 10 µL of 800 µM ATP. The mixture was then incubated at 37 °C for 10 min, before being cooled on ice for 30 s. Detection of released inorganic phosphate was achieved by adding 50 µL BIOMOL Green reagent (Enzo Life Sciences, Cat. # BML-AK111-0250) and allowing the color to develop at room temperature for 30 min. Absorbance at 635 nm was measured, and phosphate concentrations were estimated using a standard curve. ATPase activity for each sample was normalized to that of p97^WT^ controls.

### Preparation of slide surfaces for single-molecule microscopy

Glass coverslips (No. 1.5, 24 × 50 mm²; VWR, Cat. #48393-241) were modified using a mixture of mPEG-SVA and biotin-PEG-SVA in a 10:1 ratio (Laysan Bio, Cat. #MPEG-SVA-5000-1g and #BIO-PEG-SVA-5K-100MG) as previously outlined^63^. To maintain PEG integrity, functionalized coverslips were wrapped in aluminum foil and stored in a nitrogen-filled environment for up to four weeks. Immediately before imaging, custom sample chambers were prepared by trimming approximately 2 cm from the wide end of standard micropipette tips (Thermo Fisher, Cat. #02-682-261), discarding the tapered ends, and placing the tubular sections directly onto the PEG-coated coverslips. The chamber bases were securely affixed to the glass by sealing the perimeters with epoxy adhesive (Ellsworth Adhesives, Cat. #4001).

### Single-molecule fluorescence microscopy

Single-molecule fluorescence imaging was carried out on an Oxford Nanoimager (ONI) benchtop system configured for objective-type total internal reflection fluorescence (TIRF) microscopy, equipped with dual-band emission filtering and a Hamamatsu ORCA-Flash4 V3 sCMOS camera (instrument specifications: https://oni.bio/nanoimager/). A 100× oil-immersion objective (NA 1.4) was used for all measurements, and focus was stabilized using the integrated Z-lock autofocus module. Experiments were performed at ∼25 °C using the instrument’s internal temperature control; the system was allowed to equilibrate to the setpoint before acquisition, and the water-bath setting was adjusted after laser turn-on as needed to maintain temperature stability. TIRF excitation was established with an incidence angle of ∼53.5°. Cy3-labeled Ub^n^-S (n = 1–5) and Cy5-labeled proteins (^Cy5^Npl4, ^Cy5^Ufd1, and ^Cy5^p97, as indicated) were excited with 532-nm (15% power) and 640-nm (8% power) lasers, respectively (≈6 mW maximum output each), with powers chosen to maximize signal-to-noise. Unless noted otherwise, images were acquired with a 250-ms exposure time, and time series of up to 10 min were recorded per field of view.

### Single-molecule experiments for polyubiquitin substrate recognition

Single-molecule assays were conducted on biotin-PEG–passivated glass coverslips. The coverslips were washed three times with 1× T-50 buffer to ensure removal of contaminants. Streptavidin (1 mg/mL) was introduced into each sample chamber and incubated for 10 minutes to enable biotin-streptavidin interaction, followed by a rinse with 1× PBS (pH 7.4; Thermo Fisher Scientific, Cat. #10-010-049) containing 1 mg/mL BSA. Biotinylated, Cy3-labeled polyubiquitin substrates (Cy3-Ub^n^-S, n = 1–5) were added at a concentration of approximately 200 pM and incubated for 20 minutes at room temperature. Remaining unbound substrate was removed by additional washing. After substrate immobilization, chambers were refreshed with imaging buffers, and single-molecule measurements were initiated without delay.

Imaging buffer consisted of 1× PBS (pH 7.4; Gibco), 20 mM MgCl₂, 5 mM 3,4-dihydroxybenzoic acid (Fisher, #AC114891000), 0.05 mg/mL protocatechuate 3,4-dioxygenase (Sigma-Aldrich, #P8279-25UN), and 1 mM Trolox (Fisher, #218940050) to maintain an oxygen scavenging environment^64^. For experiments with ^Cy5^Npl4 alone, 20 nM ^Cy5^Npl4 was added directly to the imaging solution. To investigate Npl4**–**Ufd1 heterodimer recognition, 20 nM ^Cy5^Npl4 was first preincubated with 1 μM unlabeled Ufd1 and then Npl4**–**Ufd1 heterodimer complex was added to the imaging solution for the measurement. For Npl4**–**Ufd1**–**p97 complex formation, 20 nM ^Cy5^Npl4 was preincubated with 1 μM each of Ufd1 and p97 before chamber addition. Experiments regarding nucleotide-bound states of p97 followed the same protocol, with ATP or ATPγS added to the imaging solution to achieve the desired nucleotide condition. All assays commenced promptly after buffer exchanges to preserve sample integrity.

### Data acquisition and analysis for Cy5 labeled probe by colocalization assays

Data acquisition and analysis were performed using a custom MATLAB script^65^. The intensity-versus-time traces were extracted from the raw datasets using this code. Traces were then selected manually based on the following criteria: a single step photobleaching event of Cy3, which is labeled with Ub^n^-S, to confirm the presence of single substrate attached to the cover slip spot, as well as Cy5 fluorescence spikes exceeding five times the background intensity. Traces exhibiting binding events were subsequently idealized with a two-state model (bound and unbound) employing the segmental *k*-means algorithm implemented in QuB^65, 66^. The dwell times corresponding to the bound (τ_on_) and unbound (τ_off_) states of probe were determined from these idealized traces. Cumulative distributions of these dwell times were then generated and fit with single or double exponential functions in Origin, yielding lifetime estimates for each state. The dissociation rate constant (k_off_) was calculated as the reciprocal of τ_on_, while the association rate constant (k_on_) was determined by dividing the reciprocal of τ_off_ as well as probe concentration used during data acquisition.

### Statistical Tests

Statistical significance throughout this study was assessed using two-tailed Student’s t-tests applied to all experimental groups, ensuring a consistent and rigorous evaluation of pairwise differences in experimental outcomes. P-values were calculated based on the means and standard deviations derived from each sample group, with criteria set to accommodate the inherent biological variability observed in our assays. This process maintained methodological rigor and comparability across the various *in-vitro* scenarios examined

## Acknowledgements

We sincerely thank S. Roy (Chemistry and Biochemistry, University of Mississippi, University, Mississippi, USA) and A. Chauvier and A. Johnson-Buck (Department of Chemistry, University of Michigan, Ann Arbor, MI, USA) for their insightful discussions on the development of analysis pipelines for our single-molecule datasets. We appreciate the assistance of Emily E. Blythe and Andreas Martin (Department of Molecular and Cell Biology, University of California, Berkeley, Berkeley, CA, USA) for gifting a plasmid. We also thank Ben Dodd (Department of Human Genetics, University of Michigan, Ann Arbor, MI, USA), Andreas Schmidt (Department of Chemistry, University of Michigan, Ann Arbor, MI, USA), Jenn Russ and Ryan D. Baldridge (Department of Biological Chemistry, University of Michigan Medical School, Ann Arbor, MI, USA) for invaluable discussions related to parts of this project. N.G.W. and S.L.M. acknowledge funding from Chan Zuckerberg Initiative (CZI) collaborative pairs grants (2022-250725 and 2022-250629, respectively), and NIH grant R35 GM131922 to N.G.W.

## Author contributions

L.K. was responsible for designing the study, conducting experiments, performing data analysis, developing methodologies and software, generating visualizations, and preparing the initial manuscript draft. J.T. contributed to study design, experimental work, data interpretation, methodological and software development, visualization, and drafted the original manuscript. S.L.M. conceived the project, and contributed to methodology and experimental design, administration, funding, interpretation of results, and writing and editing the manuscript. N.G.W. oversaw project design and administration, secured funding, guided methodological approaches, supervised the research, and finalized the manuscript.

## Additional information

**Extended data** are available for this paper at…

**Supplementary information** The online version contains supplementary material available at…

## References

1. van den Boom, J. & Meyer, H. VCP/p97-mediated unfolding as a principle in protein homeostasis and signaling. Mol. Cell 69, 182–194 (2018).

2. Gwon, Y. et al. Ubiquitination of G3BP1 mediates stress granule disassembly in a context-specific manner. Science 372, eabf6548 (2021).

3. Anderson, D.J. et al. Targeting the AAA ATPase p97 as an approach to treat cancer through disruption of protein homeostasis. Cancer Cell 28, 653–665 (2015).

4. Ye, Y., Shibata, Y., Yun, C., Ron, D. & Rapoport, T. A. A membrane protein complex mediates retro-translocation from the ER lumen into the cytosol. Nature 429, 841–847 (2004).

5. Erzberger, J.P. & Berger, J.M. Evolutionary relationships and structural mechanisms of AAA+ proteins. Annu. Rev. Biophys. Biomol. Struct. 35, 93–114 (2006).

6. Brandman, O. & Hegde, R.S. Ribosome-associated protein quality control. Nat. Struct. Mol. Biol. 23, 7–15 (2016).

7. Meyer, H. & Weihl, C.C. The VCP/p97 system at a glance: connecting cellular function to disease pathogenesis. J. Cell. Sci. 127, 3877–3883 (2014).

8. Maxwell, B. A. et al. Ubiquitination is essential for recovery of cellular activities after heat shock. Science 372, eabc3593 (2021).

9. Lemberg, M.K. & Strisovsky, K. Maintenance of organellar protein homeostasis by ER-associated degradation and related mechanisms. Mol. Cell. 81, 2507–2519 (2021).

10. Wrobel, L., Hoffmann, J.L., Li, X.Y. & Rubinsztein, D.C. p37 regulates VCP/p97 shuttling and functions in the nucleus and cytosol. Sci. Adv. 10 (2024).

11. Buchan, J.R., Kolaitis, R.M., Taylor, J.P. & Parker, R. Eukaryotic stress granules are cleared by autophagy and Cdc48/VCP function. Cell 153, 1461–1474 (2013).

12. Deshaies, R.J. Proteotoxic crisis, the ubiquitin-proteasome system, and cancer therapy. Bmc Biol. 12, 94 (2014).

13. Kiss, L., James, L.C. & Schulman, B.A. UbiREAD deciphers proteasomal degradation code of homotypic and branched K48 and K63 ubiquitin chains. Mol. Cell 85, 1467–1476 (2025).

14. Williams, C., Dong, K.C., Arkinson, C. & Martin, A. The Ufd1 cofactor determines the linkage specificity of polyubiquitin chain engagement by the AAA+ ATPase Cdc48. Mol. Cell 83, 759–769 (2023).

15. Al-Obeidi, E. et al. Genotype-phenotype study in patients with valosin-containing proteinmutations associated with multisystem proteinopathy. Clin. Genet. 93, 119–125 (2018).

16. Watts, G. D. et al. Inclusion body myopathy associated with Paget disease of bone and frontotemporal dementia is caused by mutant valosin-containing protein. Nat. Genet. 36, 377–381 (2004).

17. Hübbers, C.U. et al. Pathological consequences of VCP mutations on human striated muscle. Brain 130, 381–393 (2007).

18. Custer, S. K., Neumann, M., Lu, H., Wright, A. C. & Taylor, J. P. Transgenic mice expressing mutant forms VCP/p97 recapitulate the full spectrum of IBMPFD including degeneration in muscle, brain and bone. Hum. Mol. Genet. 19, 1741–1755 (2010).

19. Blythe, E. E., Gates, S. N., Deshaies, R. J. & Martin, A. Multisystem proteinopathy mutations in VCP/p97 increase NPLOC4·UFD1L binding and substrate processing. Structure 27, 1820–1829 (2019)

20. Niwa, H., et al. The role of the N-domain in the ATPase activity of the mammalian AAA ATPase p97/VCP. J. Biol. Chem. 287, 8561–8570 (2012).

21. Blythe, E. E., Olson, K. C., Chau, V. & Deshaies, R. J. Ubiquitin- and ATP-dependent unfoldase activity of p97/VCP–NPLOC4–UFD1L is enhanced by a mutation that causes multisystem proteinopathy. Proc. Natl Acad. Sci. USA 114, E4380–E4388 (2017).

22. Banerjee, S. et al. 2.3Å resolution cryo-EM structure of human p97 and mechanism of allosteric inhibition. Science 351, 871–875 (2016).

23. Bebeacua, C. et al. Distinct conformations of the protein complex p97-Ufd1-Npl4 revealed by electron cryomicroscopy. Proc. Natl. Acad. Sci. USA 109, 1098–1103 (2012).

24. Kracht, M. et al. The accessory adapters FAF1, FAF2, and UBXN7 accelerate proteasomal degradation by increasing prior p97-mediated substrate unfolding. Sci Adv 12 (2026).

25. Bruderer, R.M., Brasseur, C. & Meyer, H.H. The AAA ATPase p97/VCP interacts with its alternative co-factors, Ufd1-Npl4 and p47, through a common bipartite binding mechanism. J. Biol. Chem. 279, 49609–49616 (2004).

26. Twomey, E. C. et al. Substrate processing by the Cdc48 ATPase complex is initiated by ubiquitin unfolding. Science 365, eaax1033 (2019).

27. Liao, Z., Arkinson, C. & Martin, A. Faf1 accelerates p97-mediated protein unfolding by promoting ubiquitin engagement. bioRxiv (2025).

28. Pan, M. et al. Seesaw conformations of Npl4 in the human p97 complex and the inhibitory mechanism of a disulfiram derivative. Nat. Commun. 12, 121 (2021).

29. Bodnar, N. O. & Rapoport, T. A. Molecular mechanism of substrate processing by the Cdc48 ATPase complex. Cell 169, 722–735 (2017).

30. Sato, Y. et al. Structural insights into ubiquitin recognition and Ufd1 interaction of Npl4. Nat. Commun. 10, 5708 (2019).

31. Shein, M. et al. Characterizing ATP processing by the AAA+ protein p97 at the atomic level. Nat. Chem. 16, 363–372 (2024).

32. Bulfer, S. L., Chou, T.-F. & Arkin, M. R. p97 disease mutations modulate nucleotide-induced conformation to alter protein-protein interactions. ACS Chem. Biol. 11, 2112–2116 (2016).

33. Huang, R., Ripstein, Z. A., Rubinstein, J. L. & Kay, L. E. Cooperative subunit dynamics modulate p97 function. Proc. Natl. Acad. Sci. USA 116, 158–167 (2019).

34. Tang, W. K. & Xia, D. Altered intersubunit communication is the molecular basis for functional defects of pathogenic p97 mutants. J. Biol. Chem. 288, 36624–36635 (2013).

35. Schuetz, A. K. & Kay, L. E. A dynamic molecular basis for malfunction in disease mutants of p97/VCP. eLife 5, e20143 (2016).

36. Rydzek, S., Shein, M., Bielytskyi, P. & Schütz, A. K. Observation of a transient reaction intermediate illuminates the mechanochemical cycle of the AAA-ATPase p97. J. Am. Chem. Soc.142, 14472–14480 (2020).

37. Wang, B. et al. Structure and ubiquitin interactions of the conserved zinc finger domain of Npl4. J. Biol. Chem. 278, 20225–20234 (2003).

38. Wagner, K. et al. Induced proximity to PML protects TDP-43 from aggregation via SUMO–ubiquitin networks. Nat. Chem. Biol. 21, 1408–1419 (2025).

39. Okatsu, K. et al. Adaptor-specific peptide inhibitors of the ubiquitin-chain-dependent unfolding activity of the human p97(VCP)-UFD1-NPL4 complex. J. Med. Chem. 68, 11270–11278 (2025).

40. Van der Meer, P. J. et al. Hierarchical mechanisms control the clearance of DNA lesion-stalled RNA polymerase II. Nat. Commun. 17, 1647 (2026).

41. Chou, T. F. et al. Specific inhibition of p97/VCP ATPase and kinetic analysis demonstrate interaction between D1 and D2 ATPase domains. J. Mol. Biol. 426, 2886–2899 (2014).

42. Tang, W. K. & Xia, D. Mutations in the human AAA+ chaperone p97 and related diseases. Front. Mol. Biosci. 3, 79 (2016).

43. Zhang, X., et al. Altered cofactor regulation with disease-associated p97/VCP mutations. Proc. Natl Acad. Sci. USA 112, E1705–E1714 (2015).

44. Schuller, J. M., Beck, F., Lössl, P., Heck, A. J. R. & Förster, F. Nucleotide-dependent conformational changes of the AAA+ ATPase p97 revisited. FEBS Lett. 590, 595–604 (2016).

45. Timachi, M. H. et al. Exploring conformational equilibria of a heterodimeric ABC transporter. Elife 6, e20236 (2017).

46. Paul, T. et al. Mechanistic insights into direct DNA and RNA strand transfer and dynamic protein exchange of SSB and RPA. Nucleic Acids Res. 53 (2025).

47. Wrobel, L. et al. Compounds activating VCP D1 ATPase enhance both autophagic and proteasomal neurotoxic protein clearance. Nat. Commun. 13, 4146 (2022).

48. Tang, W. K. et al. A novel ATP-dependent conformation in p97 N-D1 fragment revealed by crystal structures of disease-related mutants. EMBO J. 29, 2217–2229 (2010).

49. Caffrey, B. et al. AAA+ ATPase p97/VCP mutants and inhibitor binding disrupt inter-domain coupling and subsequent allosteric activation. J. Biol. Chem. 297, 101187 (2021).

50. Tong, Z.B. et al. Structural basis for E4 enzyme Ufd2-catalyzed K48/K29 branched ubiquitin chains. Nat. Chem. Biol. 22 (2026).

51. Alderson, T. R. & Kay, L. E. NMR spectroscopy captures the essential role of dynamics in regulating biomolecular function. Cell 184, 577–595 (2021).

52. Petrovic, S., et al. Structural remodeling of AAA+ ATPase p97 by adaptor protein ASPL facilitates posttranslational methylation by METTL21D. Proc. Natl Acad. Sci. USA 120, e2208941120 (2023).

53. Bard, J. A. M., Bashore, C., Dong, K. C. & Martin, A. The 26S proteasome utilizes a kinetic gateway to prioritize substrate degradation. Cell 177, 286–298 (2019).

54. Olivares, A. O., Baker, T. A. & Sauer, R. T. Mechanical protein unfolding and degradation. Annu. Rev. Physiol. 80, 413–429 (2018).

55. Tang, W.K. & Xia, D. Structural and functional deviations in disease-associated p97 mutants. J Struct Biol 179, 83–92 (2012).

56. Dong, K. C. et al. Preparation of distinct ubiquitin chain reagents of high purity and yield. Structure 19, 1053–1063 (2011).

57. Rao, M. V., Williams, D. R., Cocklin, S. & Loll, P. J. Interaction between the AAA+ ATPase p97 and its cofactor ataxin3 in health and disease: Nucleotide-induced conformational changes regulate cofactor binding. J. Biol. Chem. 292, 18392–18407 (2017).

58. Demidov, V.M., Gonchar, I.V., Tripathy, S.K., Ataullakhanov, F.I. & Grishchuk, E.L. Ndc80 complex, a conserved coupler for kinetochore-microtubule motility, is a sliding molecular clutch. Sci Adv 11 (2025).

59. Eskilson, O. et al. Nanocellulose wound dressings with integrated protease sensors for detection of wound pathogens. ACS Sens. 10, 3953–3963 (2025).

60. Xing, W.J., Li, D.Y., Wang, W.J., Liu, J.J.G. & Chen, C.L. Conformational dynamics of CasX (Cas12e) in mediating DNA cleavage revealed by single-molecule FRET. Nucleic Acids Res 52, 9014–9027 (2024).

61. Nandi, P. et al. Mechanism of allosteric inhibition of human p97/VCP ATPase and its disease mutant by triazole inhibitors. Commun. Chem. 7 (2024).

62. Green, N. et al. Antitumor Efficacy of 1,2,4-Triazole-Based VCP/p97 Allosteric Inhibitors. J. Med. Chem. 68, 14465–14494 (2025).

63. Johnson-Buck, A. et al. Kinetic fingerprinting to identify and count single nucleic acids. Nat. Biotechnol. 33, 730–732 (2015).

64. Aitken, C.E., Marshall, R.A. & Puglisi, J.D. An oxygen scavenging system for improvement of dye stability in single-molecule fluorescence experiments. Biophys. J. 94, 1826–1835 (2008).

65. Suddala, K.C. & Walter, N.G. Riboswitch Structure and Dynamics by smFRET Microscopy. Methods Enzymol. 549, 343–373 (2014).

66. Chauvier, A., Dandpat, S.S., Romero, R. & Walter, N.G. A nascent riboswitch helix orchestrates robust transcriptional regulation through signal integration (vol 15, 3955, 2024). Nat. Commun. 16 (2025).

