## Supplementary material for "Npl4 decodes polyubiquitin length and gates D1-D2 coupling in human VCP/p97": 11 Supplementary Figures and 2 Supplementary Tables

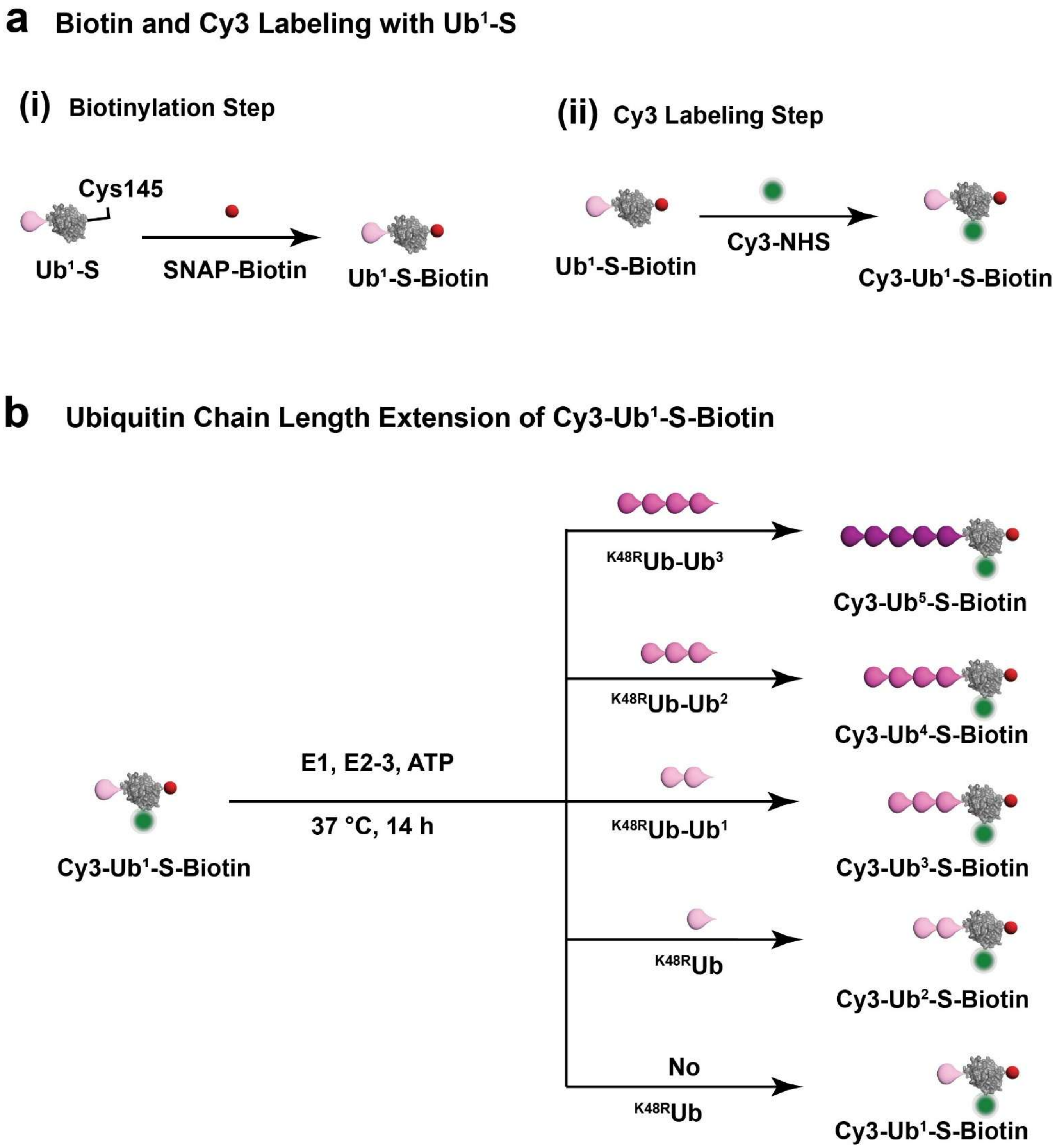


**Supplementary Fig. 1| Stepwise biotinylation, fluorescent labelling, and polyubiquitination workflow. a** Schematic overview of the sequential modification of Ub1-S, including (i) biotinylation for coverslip immobilization and (ii) Cy3 labeling for fluorescence imaging. **b** Workflow of the enzymatic polyubiquitination reaction used to generate ubiquitin substrates (here, SNAP-tag) of defined chain lengths for subsequent assays. Related to Figures 1 and 2.

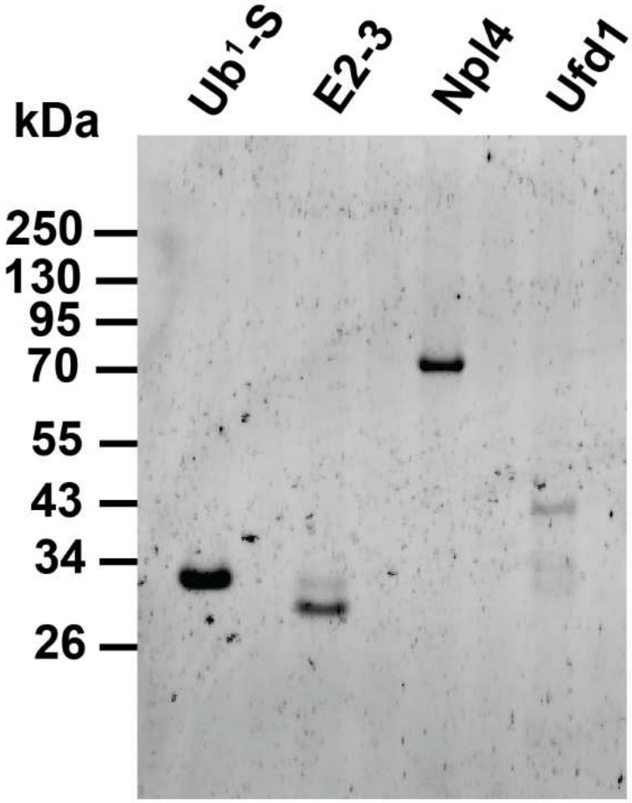


**Supplementary Fig. 2| SDS-PAGE analysis of Ub$^{1}$-S, E2-3, Npl4, and Ufd1 used in this study**.

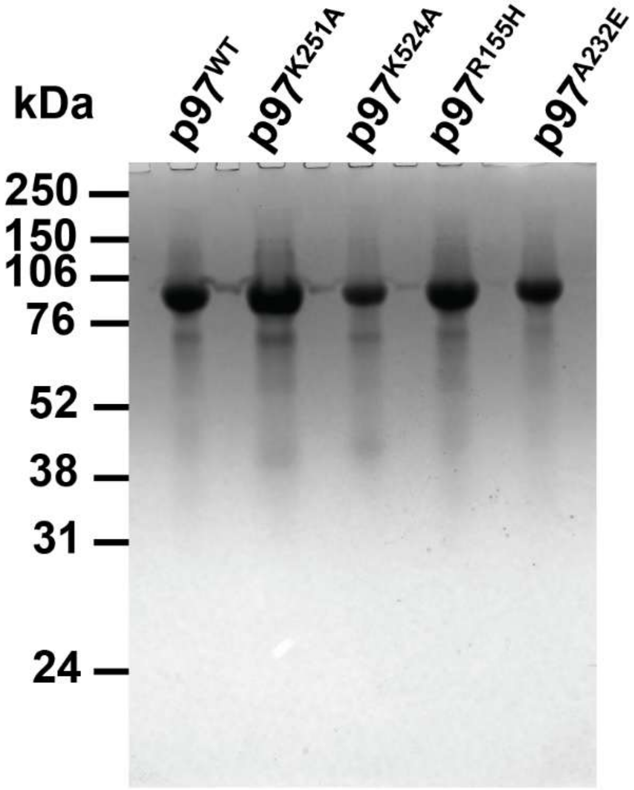


**Supplementary Fig. 3| SDS-PAGE analysis of p97 variants used in this study**.

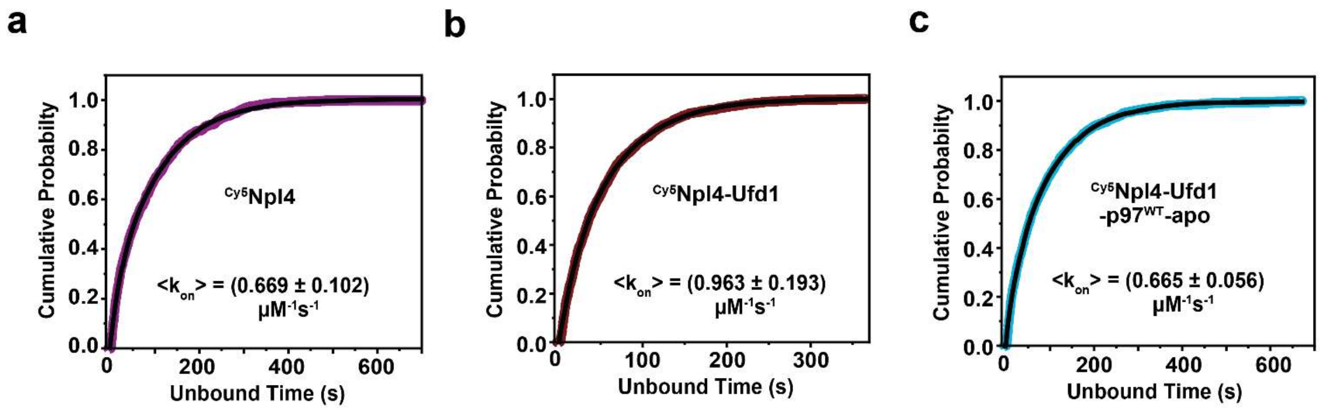


**Supplementary Fig. 4| Binding kinetics of Npl4 to Ub$^5$-S and modulation by Ufd1 and p97. a–c** Cumulative distributions of unbound dwell times for Cy5-labeled Npl4 interacting with Ub$^5$-S substrates under different conditions: (**a**) Npl4 alone, (**b**) in the formation of heterodimer with Ufd1, and (**c**) in association with the Ufd1–p97$^{WT}$ complex. Related to Figure 3

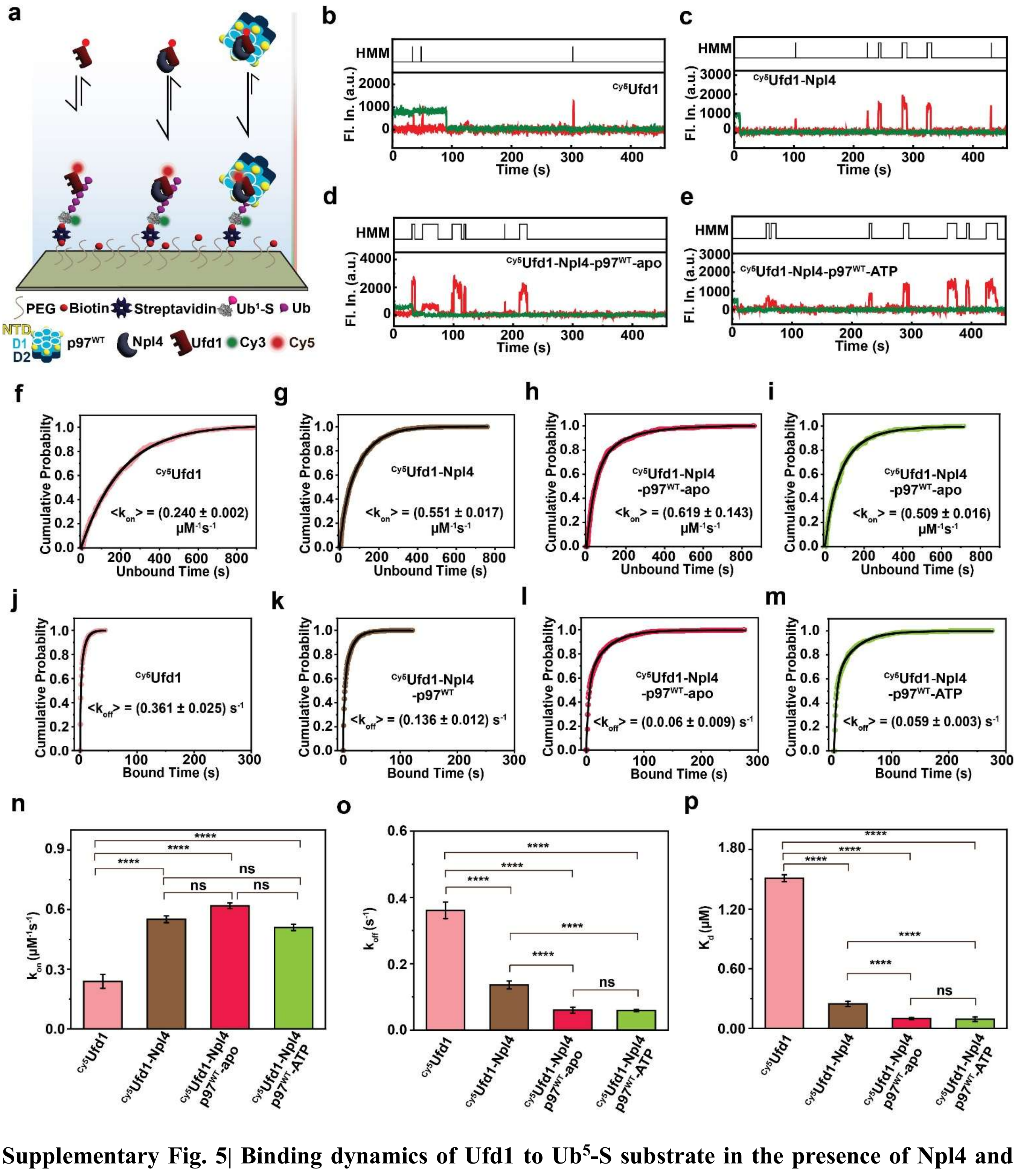


**Supplementary Fig. 5| Binding dynamics of Ufd1 to Ub$^{5}$-S substrate in the presence of Npl4 and p97$^{WT}$. a** Schematic representation of the experimental workflow used to measure association and dissociation kinetics of Cy5-labelled Ufd1 ($^{Cy5}$Ufd1) interacting with Ub$^{5}$-S substrates under three conditions: Ufd1 alone, Ufd1 following Npl4 heterodimer formation, and the Npl4–p97$^{WT}$ complex. **b–e** Representative single-molecule fluorescence intensity time traces for $^{Cy5}$Ufd1 binding to Ub$^{5}$-S under four conditions: Ufd1 alone (**b**), following assembly with Npl4 (**c**), association with the Npl4–p97$^{WT}$ complex in the absence of ATP (**d**), and after ATP addition (**e**). Segmentation via HMM analysis is indicated above each trace. **f–i** Cumulative distributions of unbound dwell times for $^{Cy5}$Ufd1 on Ub$^5$-S, contrasting conditions with or without Npl4 and the Npl4–p97$^{WT}$ complex, as shown. **j–m** Cumulative distributions of bound dwell times for $^{Cy5}$Ufd1 on Ub$^5$-S, similarly comparing the relevant conditions, as discussed earlier. **n–p** Quantification of kinetic parameters: association rate constant ($k_{on}$), dissociation rate constant ($k_{off}$), and equilibrium dissociation constant ($K_d$) for $^{Cy5}$Ufd1 binding to Ub$^5$-S under each condition. Statistical significance was determined using a two-tailed Student's t-test (ns, not significant; $0.05 < p \leq 0.5$; *$0.01 < p \leq 0.05$; **$0.001 < p \leq 0.01$; ***$0.0001 < p \leq 0.001$; ****$p \leq 0.0001$).

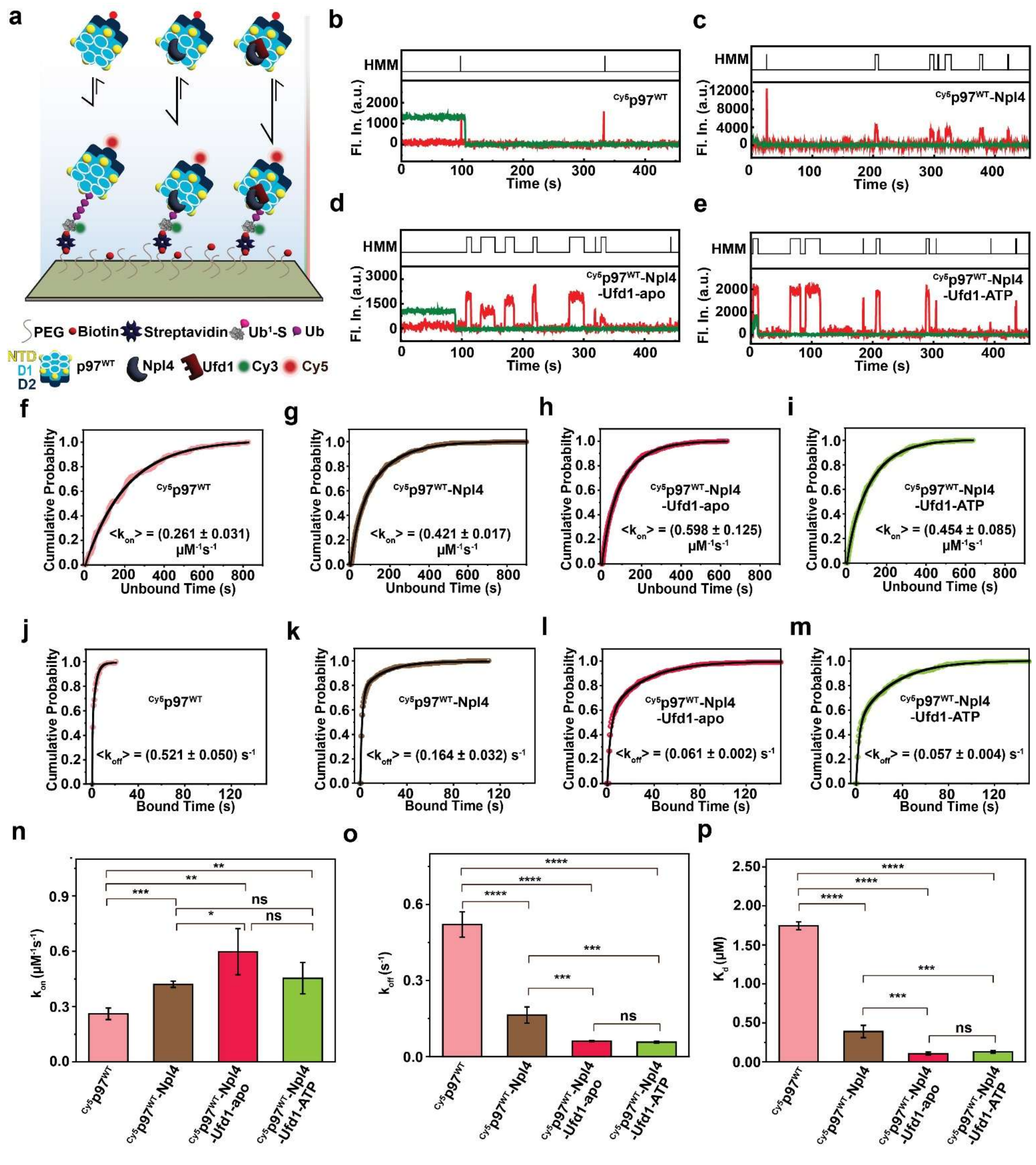


**Supplementary Fig. 6| Binding dynamics of p97$^{WT}$ to Ub$^5$-S substrate in the presence of Npl4 and Ufd1. a** Diagram outlining the experimental setup used to monitor association and dissociation kinetics of Cy5-labelled p97$^{WT}$ ($^{Cy5}$p97) binding to Ub$^5$-S substrates across three conditions: p97 alone, after incorporation with Npl4, and following assembly with the Npl4–Ufd1 complex. **b–e** Representative single-molecule fluorescence intensity trajectories for $^{Cy5}$p97 binding to Ub$^{5}$-S under four scenarios: p97 alone (**b**), after addition of Npl4 (**c**), association with the Npl4–Ufd1 complex under apo conditions (**d**), and upon ATP addition (**e**). HMM-based segmentations are shown above each trace. **f–i** Cumulative distribution curves of unbound dwell times for $^{Cy5}$p97 interacting with Ub$^{5}$-S, comparing the presence and absence of Npl4 and the Npl4–Ufd1 complex. **j–m** Cumulative distributions of bound dwell times for $^{Cy5}$p97 on Ub$^{5}$-S, comparing the specified conditions. **n–p** Quantitative analysis of kinetic parameters including association rate constant ($k_{on}$), dissociation rate constant ($k_{off}$), and equilibrium dissociation constant ($K_d$) for $^{Cy5}$p97 binding to Ub$^{5}$-S in each experimental condition. Statistical significance was determined using a two-tailed Student's t-test (ns, not significant; $0.05 < p \leq 0.5$; $*0.01 < p \leq 0.05$; $**0.001 < p \leq 0.01$; $***0.0001 < p \leq 0.001$; $****p \leq 0.0001$).

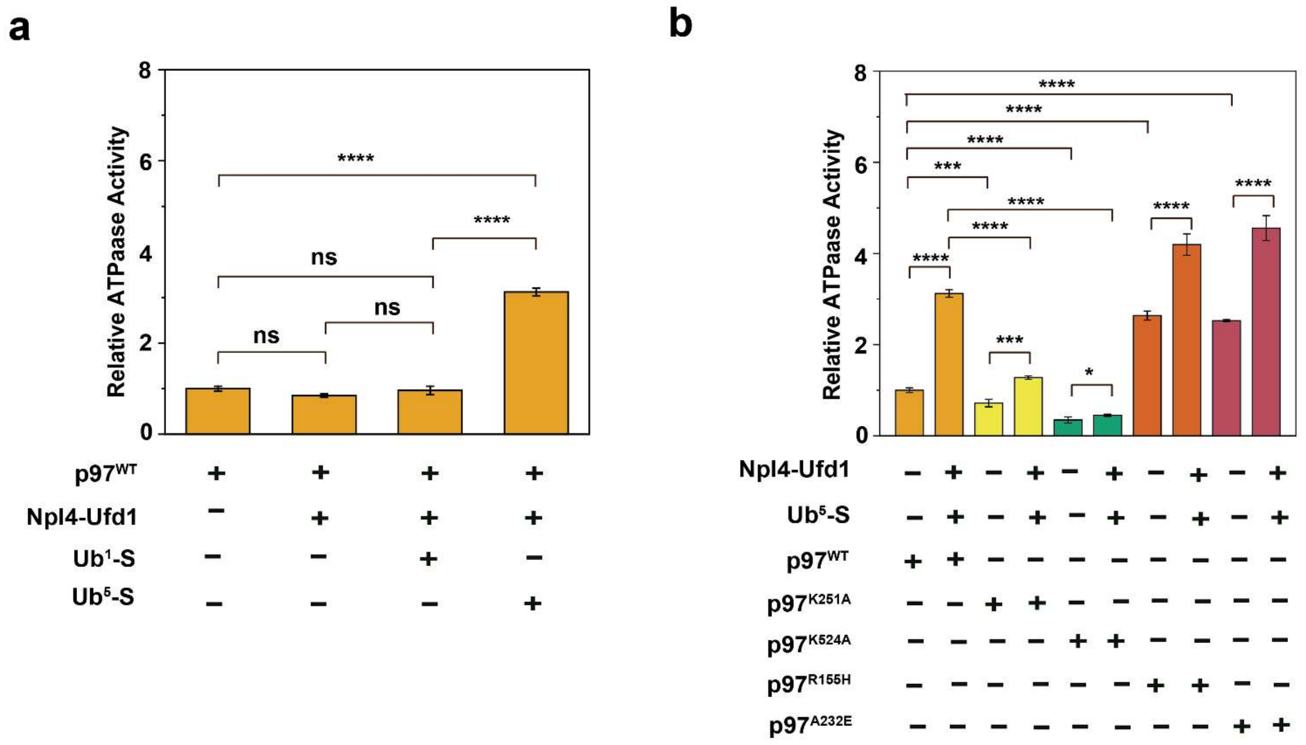


**Supplementary Fig. 7| Substrate-dependent stimulation of p97 ATPase activity in presence of UN complex. a** Relative ATPase activity of p97$^{WT}$ was assessed under various conditions: with or without the Npl4-Ufd1 cofactors, and in the presence of different ubiquitin substrates. While, Ub$^{1}$-S and UN alone did not affect p97 ATPase activity, the addition of penta-ubiquitin (Ub$^{5}$-S) resulted in a threefold increase. **b** Comparison of ATPase activity across p97 variants (WT, K251A, K524A, R155H, and A232E) with or without Npl4-Ufd1 and Ub$^{5}$-S. Measurements were performed using BIOMOL Green and normalized to basal p97$^{WT}$ activity. The D1 mutant, K251A showed a 29% reduction, and the D2 mutant K524A a 65% reduction in basal activity compared to WT. Addition of Npl4-Ufd1 and Ub$^{5}$-S led to a slight recovery in ATPase activity for these mutants. Pathogenic variants R155H and A232E exhibited elevated basal ATPase activity—4 fold and 4.5-fold above WT, respectively—consistent with previous reports. Furthermore, ATPase activity in these mutants increased significantly with the introduction of cofactors and substrate, surpassing that observed in WT and in substrate-free conditions. Data are presented as mean ± SD (n = 3). Statistical significance was determined using a two-tailed Student's t-test (ns, not significant; $0.05 < p \leq 0.5$; *$0.01 < p \leq 0.05$; **$0.001 < p \leq 0.01$; ***$0.0001 < p \leq 0.001$; ****$p \leq 0.0001$).

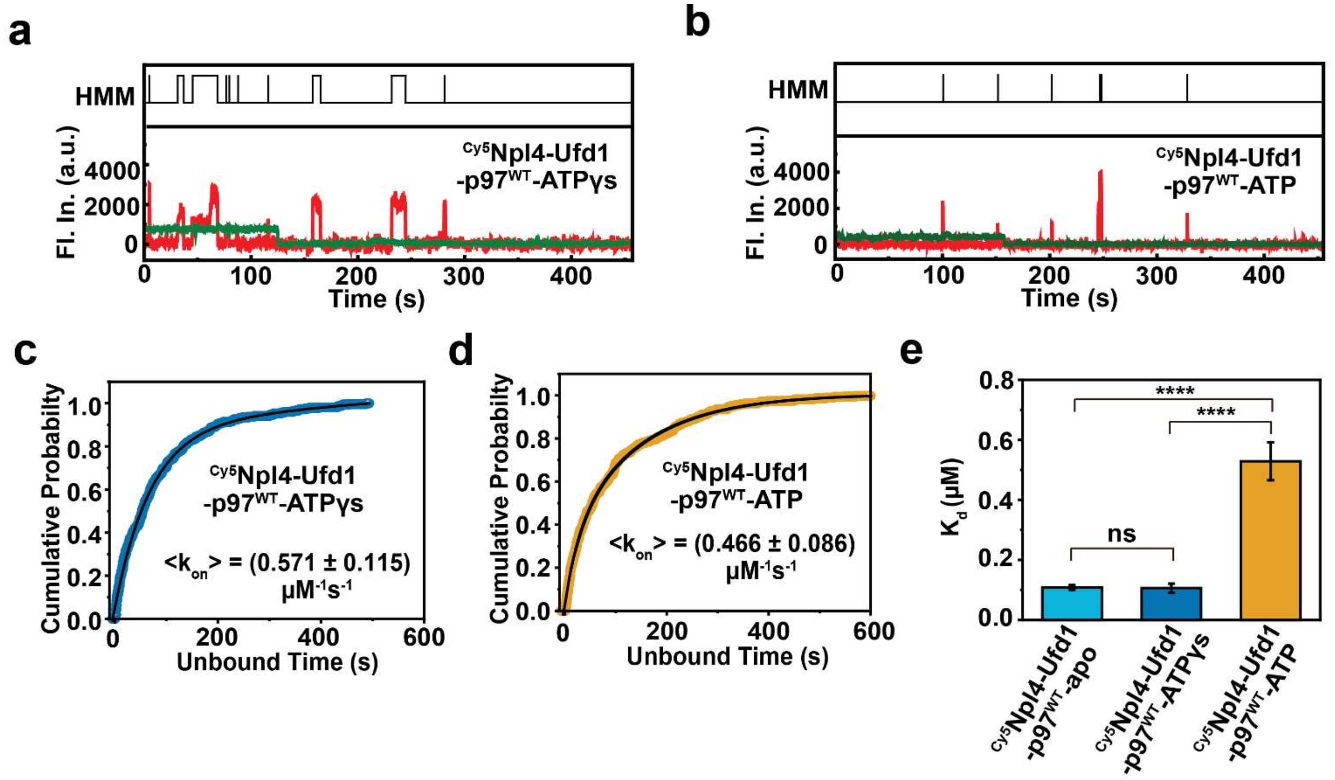


**Supplementary Fig. 8| Dynamic exchange of Npl4 within the Npl4–Ufd1–p97$^{WT}$–Ub$^{5}$-S complex is modulated by p97 conformational changes under distinct nucleotide states. a-b** Representative single-molecule fluorescence intensity trajectories for Cy5-labelled Npl4 ($^{Cy5}$Npl4) (preassembled with Npl4–Ufd1–p97$^{WT}$) bound to Ub$^{5}$-S substrate, under ATPγS (**a**) and ATP (**b**) conditions, illustrating nucleotide-dependent alterations in Npl4 binding dynamics. HMM-derived segmentations are displayed above each trace. **c**–**d** Cumulative distributions of unbound dwell times for $^{Cy5}$Npl4 (preassembled with Npl4–Ufd1–p97$^{WT}$) on Ub$^{5}$-S substrate under ATPγS (**c**) and ATP (**d**) conditions, highlighting unaltered in association rate constants related to nucleotide state. **e** Quntifiaction of equilibrium dissociation constant ($K_d$) for $^{Cy5}$Npl4 binding to Ub$^{5}$-S across the two nucleotide states. Statistical significance was evaluated by two-tailed Student's t-test (ns, not significant; $0.05 < p \leq 0.5$; *$0.01 < p \leq 0.05$; **$0.001 < p \leq 0.01$; ***$0.0001 < p \leq 0.001$; ****$p \leq 0.0001$). Related to Figure 4.

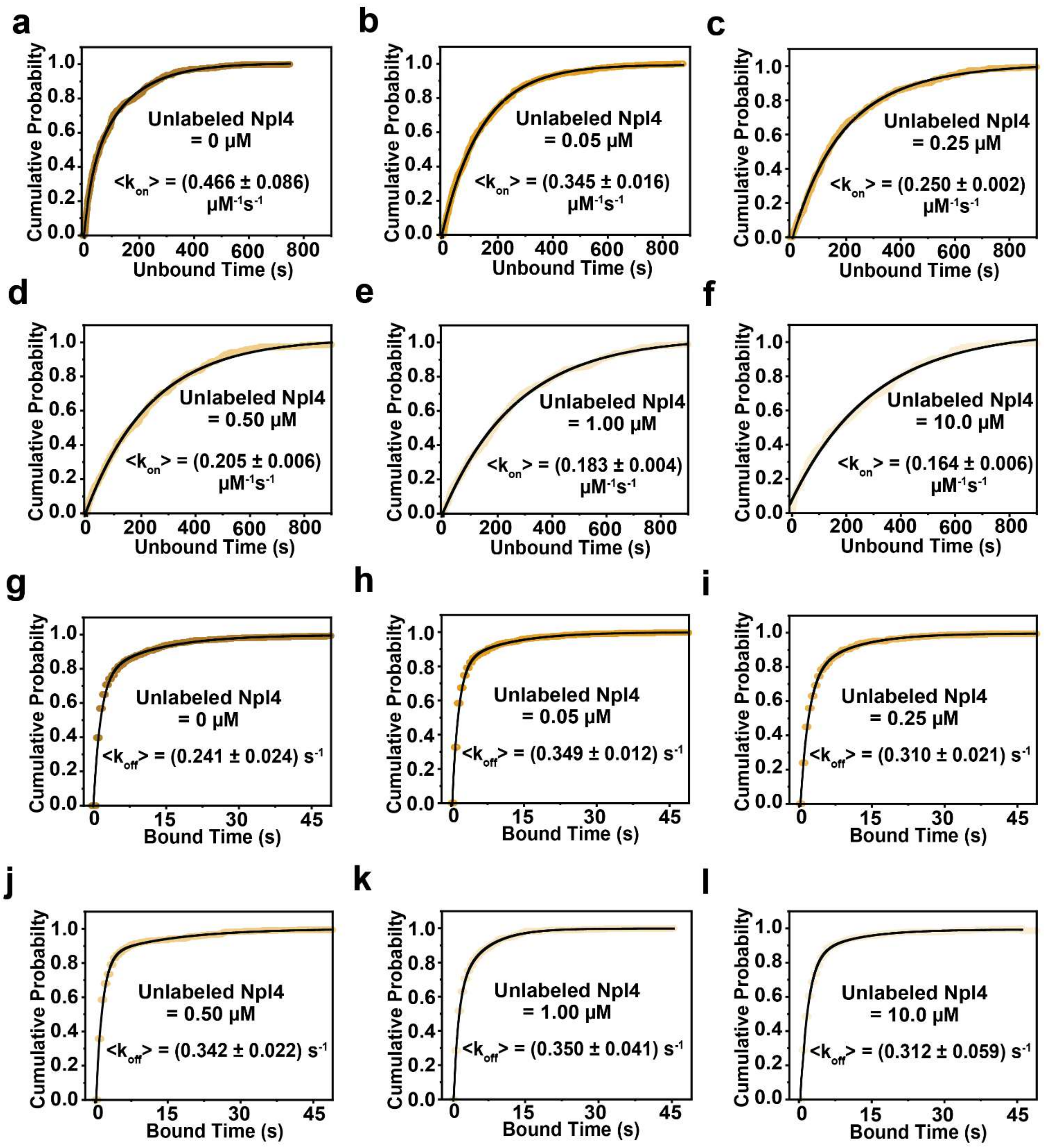


**Supplementary Fig. 9| Conformational changes in p97 during ATP hydrolysis modulate dynamic Npl4 exchange within the Npl4–Ufd1–p97$^{WT}$–Ub$^{5}$-S complex. a–f** Cumulative distributions of unbound dwell times for Cy5-labeled Npl4 ($^{Cy5}$Npl4), preassembled with the Npl4–Ufd1–p97$^{WT}$ complex, on Ub$^{5}$-S substrate under ATP hydrolysis conditions, with increasing concentrations of unlabeled Npl4 ([unlabeled Npl4] = 0, 0.05, 0.25, 0.50, 1.0, and 10.0 μM; $^{Cy5}$Npl4 = 20 nM). The data indicate a stepwise increase in association rate constants as the concentration of unlabeled Npl4 rises, reflecting enhanced Npl4 exchange within the complex during p97 ATP hydrolysis. **g–i** Cumulative distributions of bound dwell times for $^{Cy5}$Npl4 (preassembled with Npl4–Ufd1–p97$^{WT}$) bound to Ub$^{5}$-S substrate under identical ATP hydrolysis conditions and increasing concentrations of unlabeled Npl4, demonstrate that dissociation rate constants remain largely unchanged, regardless of added unlabeled Npl4. Figure 4 for related data.

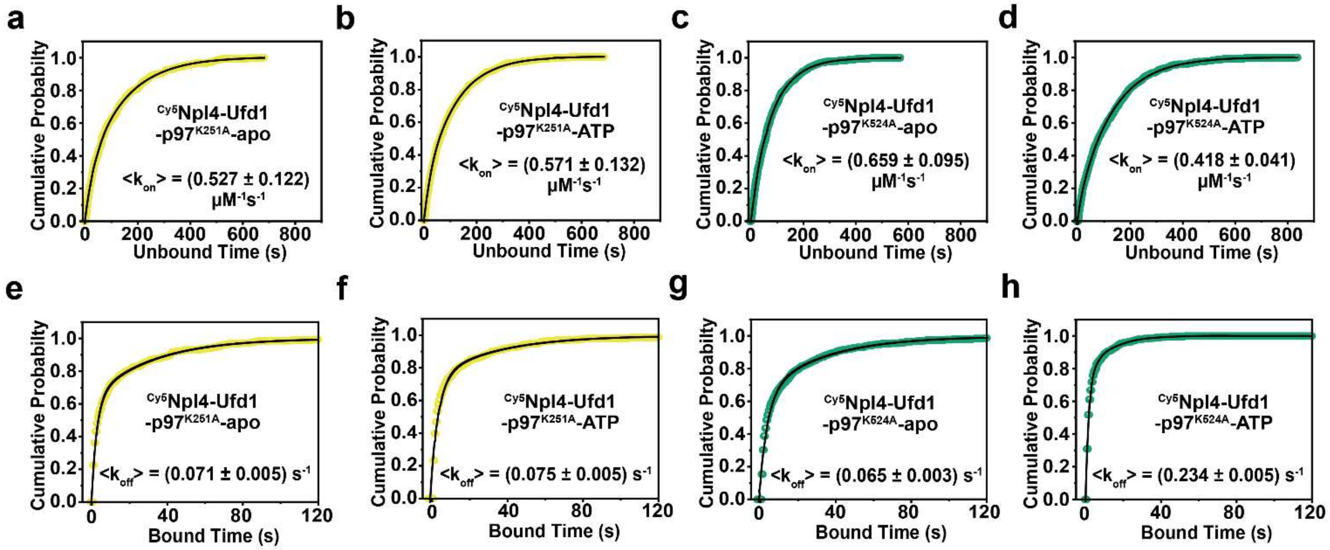


**Supplementary Fig. 10| D1-specific ATP hydrolysis facilitates dynamic Npl4 exchange on substrate.** **a-b** Cumulative distributions of unbound dwell times for $^{Cy5}$Npl4, preassembled with the Npl4–Ufd1–p97$^{K251A}$ complex (D1 domain inactive, D2 domain active), interacting with Ub$^{5}$-S substrate in the absence (**a**) and presence (**b**) of ATP. **c-d** Cumulative distributions of unbound dwell times for $^{Cy5}$Npl4 preassembled with the Npl4–Ufd1–p97$^{K524A}$ complex (D1 domain active, D2 domain inactive) on Ub$^{5}$-S substrate, without ATP (**c**) and after ATP addition (**d**). **e-f** Cumulative distributions of bound dwell times for $^{Cy5}$Npl4 with the Npl4–Ufd1–p97$^{K251A}$ complex on Ub$^{5}$-S, comparing conditions without (**e**) and with (**f**) ATP. **g-h** Cumulative distributions of bound dwell times for $^{Cy5}$Npl4 with the Npl4–Ufd1–p97$^{K524A}$ complex on Ub$^{5}$-S substrate, without (**g**) and with (**h**) ATP. See Figure 4 for related results.

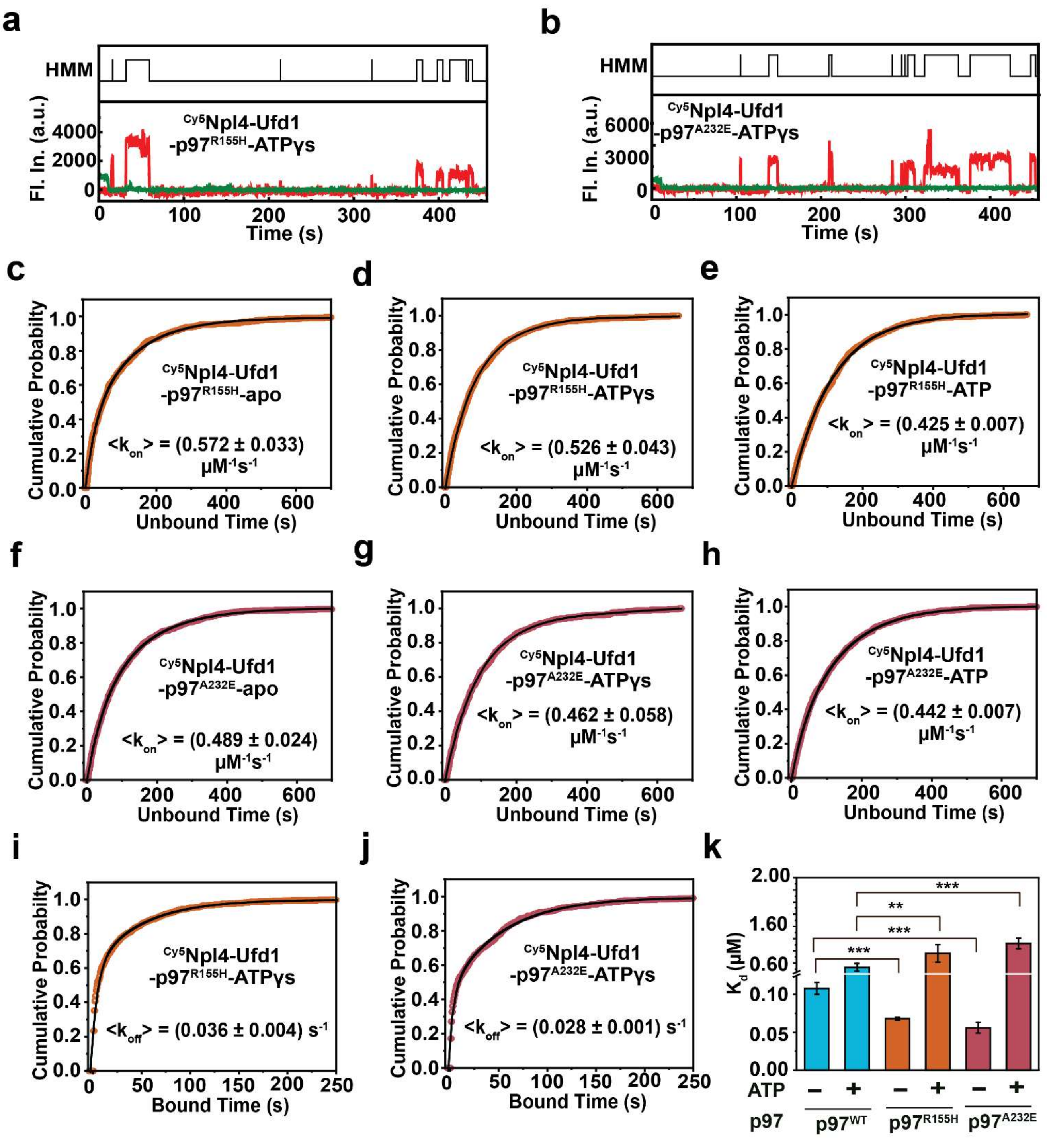


**Supplementary Fig. 11| Disease-associated p97 mutants R155H and A232E enhance substrate affinity and disrupt Npl4–Ufd1–p97 complex dynamics. a-b** Representative single-molecule fluorescence intensity trajectories for $^{Cy5}$Npl4, preassembled with Npl4–Ufd1–p97, binding to Ub$^5$-S substrate under ATPγS conditions for the R155H (**a**) and A232E (**b**) p97 variants. HMM-segmented regions are indicated above each trace. **c–e** Cumulative distributions of unbound dwell times for $^{Cy5}$Npl4 (assembled with Npl4–Ufd1–p97$^{R155H}$) binding to Ub$^5$-S substrate in the nucleotide-free, apo (**c**), ATPγS (**d**), and ATP (**e**) conditions, showing minimal changes in association rate constants across nucleotide states. **f–h** Cumulative distributions of bound dwell times for $^{Cy5}$Npl4 (with Npl4–Ufd1–p97$^{A232E}$) on Ub$^{5}$-S substrate under apo (**f**), ATPγS (**g**), and ATP (**h**) conditions, indicating similar association kinetics regardless of nucleotide state. **i** Quantification of equilibrium dissociation constant ($K_d$) for $^{Cy5}$Npl4 binding to Ub$^{5}$-S across apo and nucleotide conditions for p97$^{WT}$ and the pathogenic mutants p97$^{R155H}$ and p97$^{A232E}$. Statistical significance determined using two-tailed Student's t-test (ns, not significant; $0.05 < p \leq 0.5$; *$0.01 < p \leq 0.05$; **$0.001 < p \leq 0.01$; ***$0.0001 < p \leq 0.001$; ****$p \leq 0.0001$). See Figure 5 for related data.

**Table S1: Proteins used in this study**

| Plasmid name | Vector | Antibiotic Resistance | Expression | Purification | Comments | Source |
|---|---|---|---|---|---|---|
| Ub[1]-SNAP-tag-6xHis | pET-28a | KAN | BL21 (DE3), 0.6 mM IPTG, TB, 18 °C for 14 h | Ni-NTA, His-Trap column | - | This study |
| 6xHis-TEV-gp78-Ube2g2 | pET-28a | KAN | BL21 (DE3), 0.4 mM IPTG, TB, 25 °C for 16 h | Ni-NTA, TEV cleavage, His-Trap column | - | *Proc. Natl Acad. Sci. USA* **114**, E4380–E4388 (2017) Ref[2] |
| Npl4 | pET41b+ | AMP | BL21 (DE3), 0.4 mM IPTG, TB, 18 °C for 14 h | Ni-NTA, His-Trap column | Co-express and co-purified with Ufd1-8XHis | Addgene |
| 6xHis-TEV-Npl4 | pET-28a | KAN | BL21 (DE3), 0.4 mM IPTG, TB, 18 °C for 14 h | Ni-NTA, TEV cleavage, His-Trap column | - | This study |
| Ufd1-8XHis | pET41b+ | KAN | BL21 (DE3), 0.4 mM IPTG, TB, 18 °C for 14 h | Ni-NTA, His-Trap column | - | Addgene |
| 6xHis-TEV-Ufd1 | pET-28a | KAN | BL21 (DE3), 0.4 mM IPTG, TB, 18 °C for 14 h | Ni-NTA, TEV cleavage, His-Trap column | - | This Study |
| p97$^{WT}$-HA-6xHis | pET-28a | KAN | BL21 (DE3), 1 mM IPTG, TB, 25 °C for 12 h | Ni-NTA, His-Trap column | - | This study |

| | | | | | | |
|---|---|---|---|---|---|---|
| p97$^{K251A}$-HA-6xHis | pET-28a | KAN | BL21 (DE3), 1 mM IPTG, TB, 25 °C for 12 h | Ni-NTA, His-Trap column | - | This study |
| p97$^{K524A-}$HA-6xHis | pET-28a | KAN | BL21 (DE3), 1 mM IPTG, TB, 25 °C for 12 h | Ni-NTA, His-Trap column | - | This study |
| p97$^{R155H}$-HA-6xHis | pET-28a | KAN | BL21 (DE3), 1 mM IPTG, TB, 25 °C for 12 h | Ni-NTA, His-Trap column | - | This study |
| p97$^{A232E}$-HA-6xHis | pET-28a | KAN | BL21 (DE3), 1 mM IPTG, TB, 25 °C for 12 h | Ni-NTA, His-Trap column | - | This study |

**Table S2: Commercial proteins and chemicals used in this study**

| **Protein name** | **Catalog Number#** | **Source** |
|---|---|---|
| UBE1 | UB-0101-005 | LifeSensors |
| Ub | U-100H | Bio-Techne |
| $^{K48R}$Ub[1] | SI217 | LifeSensors |
| $^{K48R}$Ub-Ub[1] | SI4802 | LifeSensors |
| $^{K48R}$Ub-Ub[2] | SI4803 | LifeSensors |
| $^{K48R}$Ub-Ub[3] | SI4804 | LifeSensors |
| Cyanine3 NHS ester | 11020 | Lumiprobe |
| Cyanine5 NHS ester | 13020 | Lumiprobe |
